# Connectome-wide mega-analysis identifies a reproducible functional network signature of temporal lobe epilepsy

**DOI:** 10.64898/2026.03.11.711198

**Authors:** Ke Xie, Judy Chen, Ella Sahlas, Fatemeh Fadaie, Thaera Arafat, Raúl Rodríguez-Cruces, Marlo Naish, Alexander Ngo, Arielle Dascal, Alexander Barnett, Samantha Audrain, Raluca Pana, Sara Larivière, Aristides Hadjinicolaou, Alexander G. Weil, Sami Obaid, Roy Dudley, Dewi V. Schrader, Zhiqiang Zhang, Luis Concha, Andrea Bernasconi, Neda Bernasconi, Boris C. Bernhardt

## Abstract

Resting-state functional magnetic resonance imaging (MRI) studies have reported abnormal intrinsic functional connectivity (FC) across distributed circuits in patients with temporal lobe epilepsy (TLE), indicating a system-level impact of the disorder. However, findings remain inconsistent due to limited sample sizes and methodological heterogeneity, leaving the structural determinants and clinical relevance of FC alterations unresolved. To identify a robust and reproducible FC signature, we conducted a data-driven, connectome-wide mega-analysis in a large multicentre cohort of 652 participants (297 TLE, 73 disease controls, and 282 healthy controls) with multimodal 3T MRI and deep clinical phenotyping. We identified convergent FC reconfigurations at both group and individual levels that preferentially involved densely connected hubs, manifesting as hyperconnectivity in frontoparietal association systems and hypoconnectivity in temporal and paralimbic systems. Integrating structural cortical wiring features further revealed that these extensive alterations were constrained by corticocortical proximity, microstructural similarity, and white matter connectivity. Clinically, the FC phenotype tracked symptom burden and disease progression, informed postsurgical seizure outcome, and distinguished TLE from other focal epilepsies. Collectively, these findings systematically delineate a neurobiologically grounded, hub-centric pattern of intrinsic network disruption in TLE, anchored in temporolimbic and adjacent transmodal systems, with potential utility for individualized phenotypic stratification and outcome prognostication.

## 1. Introduction

Temporal lobe epilepsy (TLE) is the most common focal epilepsy in adults and is linked to pathological alterations of the mesial temporal lobe (*1*, *2*). Despite continuous advances in antiseizure medication development, approximately one third of patients with TLE remain pharmacoresistant, which is commonly associated with higher seizure burden, increased risk of seizure-related injuries and mortality, and reduced quality of life (*3*, *4*). For these individuals, surgical resection of the epileptogenic zone can offer the possibility of long-term seizure freedom (*5*). However, reliable neuroimaging biomarkers that enable the quantification of disease severity, guide presurgical evaluation and predict seizure outcome remain limited, posing a significant obstacle to precision epilepsy care (*6*).

Resting-state functional MRI (rs-fMRI) has emerged as a powerful, non-invasive technique to profile intrinsic brain function (*7-10*), capture inter-individual variability (*11-14*), and deliver clinically relevant biomarkers of dysfunction (*15-19*). Prior rs-fMRI applications in TLE have demonstrated changes in intrinsic functional connectivity (FC) across distributed cortical and subcortical circuits, supporting a network-level conceptualization of the condition (*20-23*). Despite a growing literature (*24-27*), there remains no consensus on the spatial organization, directionality, and clinical significance of FC changes in TLE. Several methodological factors contribute to these discrepancies. First, most prior studies relied on small, single-centre cohort datasets, yielding findings that do not reliably replicate across different data acquisition contexts. Second, analyses have often focused on predefined brain regions or connections of interest, or on coarse summaries of network activity, obscuring the heterogeneity of vulnerability across brain regions. Third, emphasis has mainly been placed on describing patterns at the group level, without specifically addressing inter-individual differences that may inform patient-specific stratification and treatment (*23*, *28-31*). Finally, reliance on cross-sectional study designs has precluded drawing inferences about how disease progression affects cortical dysfunction in TLE, even as evidence accumulates for progressive structural abnormalities (*32-34*).

Converging evidence from histological and neuroimaging studies of healthy individuals indicates that intrinsic FC is fundamentally shaped by the brain’s spatial embedding and structural scaffold (*35-38*). Specifically, brain regions connected by direct white matter tracts or regions in close proximity tend to exhibit stronger FC. Structural and functional connectivity gradually uncouple in transmodal association cortices, consistent with the hierarchical specialization of cortical organization (*39*). Such structure-function coupling is thought to support the coordination and propagation of neural activity across distributed regions and to shape both healthy and diseased networks (*40-43*). Developmental MRI studies show that, as large-scale cortical systems increasingly functionally differentiate during adolescence, functional coupling particularly between unimodal and transmodal systems decreases (*41*). Similarly, in patients with TLE, elevated excitation-inhibition imbalance is observed in cortical regions strongly linked to disease epicentres, highlighting network-based regional vulnerability (*44*). Still, whether, and to what extent, the brain’s structural layout accounts for the spatial heterogeneity of altered FC in TLE remains unexplored. Furthermore, the specificity of these functional changes to TLE, as opposed to their being shared across focal epilepsies, also remains unclear. Addressing these questions requires multimodal datasets and integrative analyses capable of situating large-scale FC abnormalities within the brain’s multiscale architecture.

In this study, we addressed these gaps by conducting a data-driven, connectome-wide mega-analysis of resting-state FC in one of the largest multicentre TLE cohorts examined with multimodal MRI to date, comprising 297 patients with TLE, 73 disease controls with extratemporal focal epilepsy (extra-TLE), and 282 healthy controls, all with high-resolution multimodal MRI (**Figure 1a**) (*23*). Standardized multimodal MRI processing and feature integration, data quality control, and derivative harmonization using open-access workflows were implemented across sites, and individual-level connectomes were constructed (**Figure 1b**). First, we quantified FC alterations in TLE patients across different spatial scales (edge, network, and region) at both group and individual levels using whole-brain statistical inference. Second, we examined the structural determinants of FC alterations of TLE by testing associations with structural cortical wiring using complementary indices, including cortical geometry, intracortical microstructure, and white matter connectivity. Finally, we evaluated clinical significance by relating patient-specific FC signatures to symptom severity with multivariate models and by examining associations with longitudinal changes and postsurgical outcomes (**Figure 1c**).

**Figure 1.**
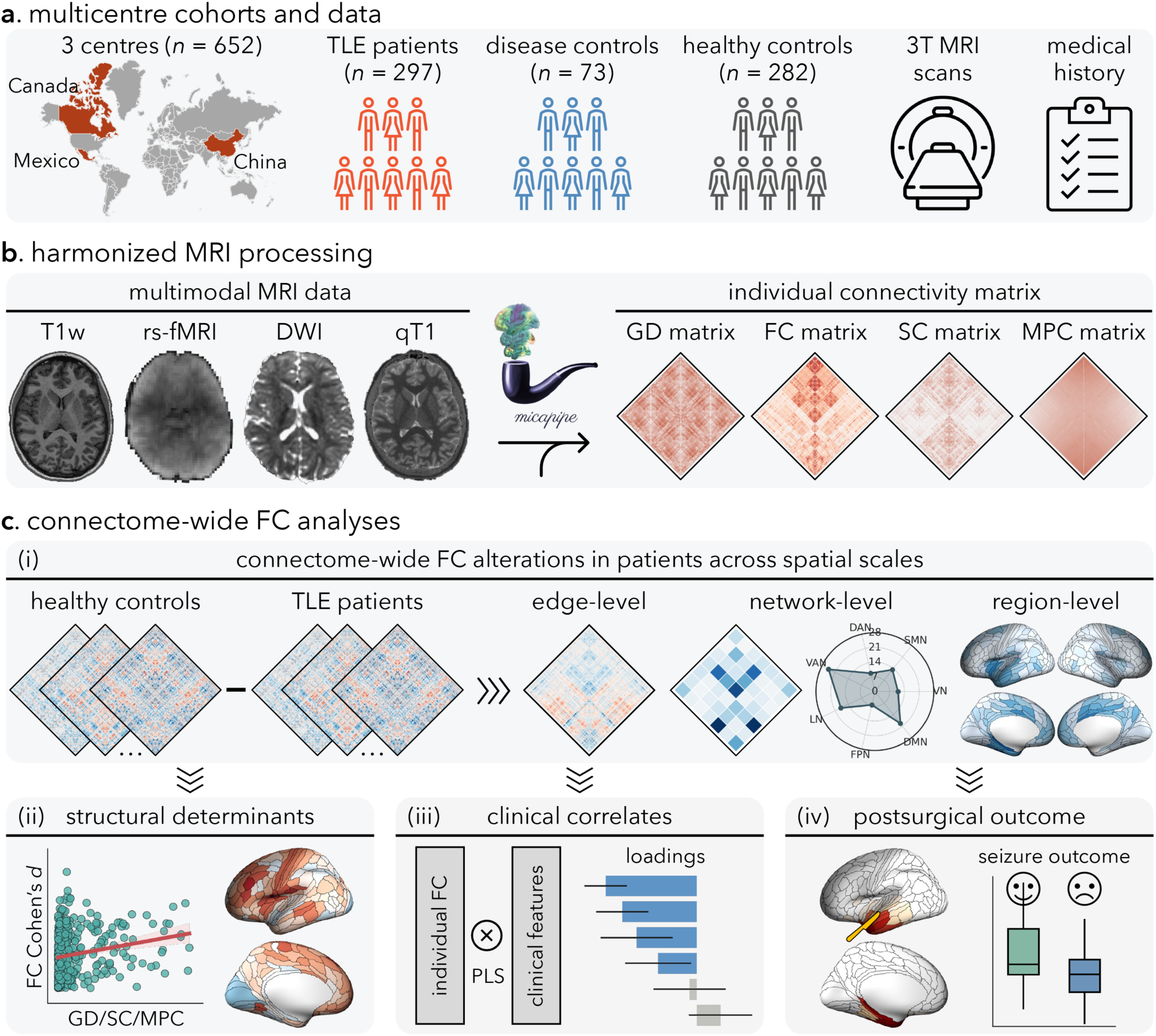
Overview of study design and analyses. **(a)** Multicentre cohorts and data. Multimodal 3T MRI and clinical data (for patient groups) were pooled across three epilepsy centres (total *n*=652), including 297 TLE patients, 73 disease controls, and 282 healthy controls. **(b)** Harmonized MRI processing. Multimodal MRI (T1w, rs-fMRI, DWI, and quantitative T1 [qT1]) were preprocessed using a standardized pipeline to generate individual connectomes: geodesic distance (GD), functional connectivity (FC), structural connectivity (SC), and microstructural profile covariance (MPC). **(c)** Connectome-wide FC analyses. **(i)** Group- and individual-level FC alterations in patients were quantified across different spatial scales (edge-, network-, and region-level). **(ii)** Structural determinants of FC alterations were determined by relating effect sizes (FC Cohen’s *d*) to cortical wiring features (GD, SC, and MPC). **(iii)** Clinical correlates were evaluated using multivariate analysis (partial least squares; PLS) linking individual FC patterns to clinical features. **(iv)** Prognostic utility of FC was assessed by testing associations between individual FC alterations and postsurgical seizure outcome.

## 2. Results

### 2.1 Participants

This multicentre, retrospective study included 652 participants: 297 patients with unilateral TLE, 73 disease controls with unilateral extra-TLE focal epilepsy, and 282 healthy controls. Participants were aggregated from four independent datasets across three centres: (i) Montreal Neurological Institute– Hospital (MICA-MICs: 72 TLE, 45 disease controls, 100 healthy controls; NOEL: 72 TLE, 28 disease controls, 42 healthy controls) (*45*), (ii) Universidad Nacional Autónoma de México (EpiC: 29 TLE, 34 healthy controls) (*46*), and (iii) Jinling Hospital, Nanjing University School of Medicine (Nanj: 124 TLE, 106 healthy controls). Details on participant inclusion and exclusion criteria are provided in the **Methods** and in our prior work (*23*). Demographic and clinical information for participants in each dataset is shown in **Table 1**. The overview of the study design and data processing is shown in **Figure 1**.

**Table 1.**
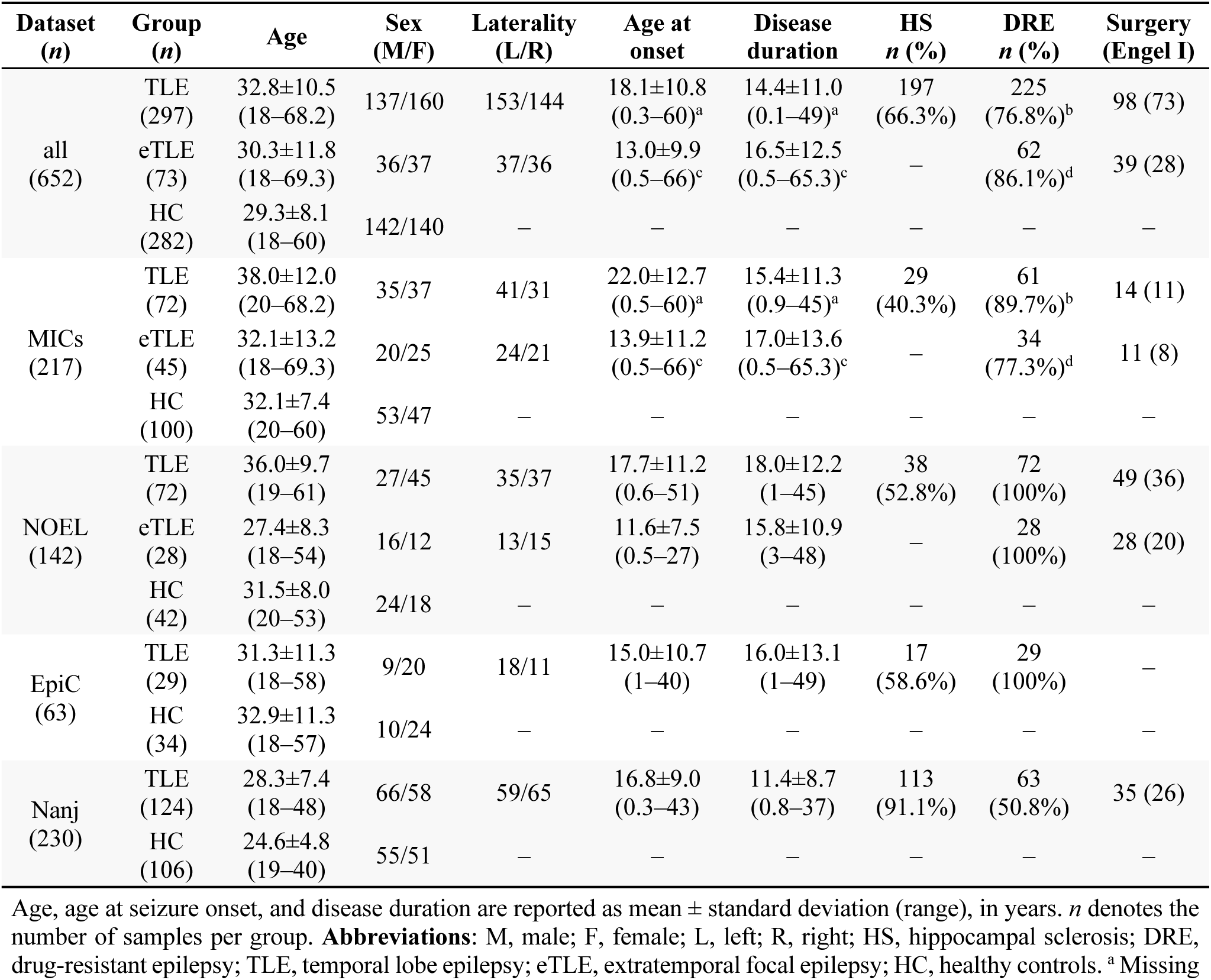
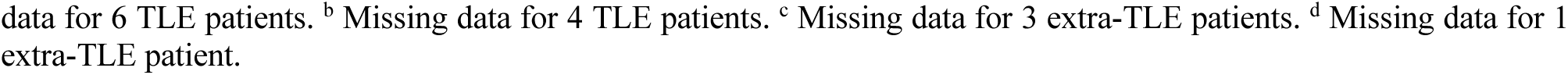
Demographic and clinical information.

| Dataset<br>(*n*) | Group<br>(*n*) | Age | Sex<br>(M/F) | Laterality<br>(L/R) | Age at<br>onset | Disease<br>duration | HS<br>*n* (%) | DRE<br>*n* (%) | Surgery<br>(Engel I) |
| --- | --- | --- | --- | --- | --- | --- | --- | --- | --- |
| all<br>(652) | TLE<br>(297) | 32.8±10.5<br>(18–68.2) | 137/160 | 153/144 | 18.1±10.8<br>(0.3–60)<sup>a</sup> | 14.4±11.0<br>(0.1–49)<sup>a</sup> | 197<br>(66.3%) | 225<br>(76.8%)<sup>b</sup> | 98 (73) |
|  | eTLE<br>(73) | 30.3±11.8<br>(18–69.3) | 36/37 | 37/36 | 13.0±9.9<br>(0.5–66)<sup>c</sup> | 16.5±12.5<br>(0.5–65.3)<sup>c</sup> | – | 62<br>(86.1%)<sup>d</sup> | 39 (28) |
|  | HC<br>(282) | 29.3±8.1<br>(18–60) | 142/140 | – | – | – | – | – | – |
| MICs<br>(217) | TLE<br>(72) | 38.0±12.0<br>(20–68.2) | 35/37 | 41/31 | 22.0±12.7<br>(0.5–60)<sup>a</sup> | 15.4±11.3<br>(0.9–45)<sup>a</sup> | 29<br>(40.3%) | 61<br>(89.7%)<sup>b</sup> | 14 (11) |
|  | eTLE<br>(45) | 32.1±13.2<br>(18–69.3) | 20/25 | 24/21 | 13.9±11.2<br>(0.5–66)<sup>c</sup> | 17.0±13.6<br>(0.5–65.3)<sup>c</sup> | – | 34<br>(77.3%)<sup>d</sup> | 11 (8) |
|  | HC<br>(100) | 32.1±7.4<br>(20–60) | 53/47 | – | – | – | – | – | – |
| NOEL<br>(142) | TLE<br>(72) | 36.0±9.7<br>(19–61) | 27/45 | 35/37 | 17.7±11.2<br>(0.6–51) | 18.0±12.2<br>(1–45) | 38<br>(52.8%) | 72<br>(100%) | 49 (36) |
|  | eTLE<br>(28) | 27.4±8.3<br>(18–54) | 16/12 | 13/15 | 11.6±7.5<br>(0.5–27) | 15.8±10.9<br>(3–48) | – | 28<br>(100%) | 28 (20) |
|  | HC<br>(42) | 31.5±8.0<br>(20–53) | 24/18 | – | – | – | – | – | – |
| EpiC<br>(63) | TLE<br>(29) | 31.3±11.3<br>(18–58) | 9/20 | 18/11 | 15.0±10.7<br>(1–40) | 16.0±13.1<br>(1–49) | 17<br>(58.6%) | 29<br>(100%) | – |
|  | HC<br>(34) | 32.9±11.3<br>(18–57) | 10/24 | – | – | – | – | – | – |
| Nanj<br>(230) | TLE<br>(124) | 28.3±7.4<br>(18–48) | 66/58 | 59/65 | 16.8±9.0<br>(0.3–43) | 11.4±8.7<br>(0.8–37) | 113<br>(91.1%) | 63<br>(50.8%) | 35 (26) |
|  | HC<br>(106) | 24.6±4.8<br>(19–40) | 55/51 | – | – | – | – | – | – |

Age, age at seizure onset, and disease duration are reported as mean ± standard deviation (range), in years. *n* denotes the number of samples per group. **Abbreviations:** M, male; F, female; L, left; R, right; HS, hippocampal sclerosis; DRE, drug-resistant epilepsy; TLE, temporal lobe epilepsy; eTLE, extratemporal focal epilepsy; HC, healthy controls. <sup>a</sup> Missing
data for 6 TLE patients. <sup>b</sup> Missing data for 4 TLE patients. <sup>c</sup> Missing data for 3 extra-TLE patients. <sup>d</sup> Missing data for 1 extra-TLE patient.

### 2.2 Functional Connectivity Alterations in TLE

#### 2.2.1 Group-Level FC Alterations

Surface-based time series for each participant were extracted from processed and denoised rs-fMRI data across 360 cortical brain regions defined by the HCP-MMP1.0 atlas (*47*). A cortex-wide FC matrix (360×360) containing 64,620 unique edges was constructed by calculating Pearson’s correlation coefficients between time series. Edge-wise between-group analysis of FC found that patients with TLE exhibited both increased FC (hyperconnectivity) and decreased FC (hypoconnectivity) compared to healthy controls (**Figure 2a**). The TLE–control FC difference matrix negatively correlated with the mean FC matrix of healthy individuals (*r*=−0.42, *P*<0.001). That is, strongly connected edges in the healthy brain demonstrated the most significant FC reductions in patients, whereas weakly connected edges showed relative FC elevation, indicating that TLE pathophysiology preferentially disrupts hub-to-hub connections that form the core of large-scale communication. At the regional level, the effect size of FC alteration was significantly correlated with regional weighted degree centrality (*rho*=−0.45, *P*_spin_<0.001; **Supplementary Figure 1**), implying greater FC reductions in high-degree hubs. At the edge level, we observed significant hyperconnectivity in 2,135 edges (3.3% of all edges; mean±SD Cohen’s *d*=0.30±0.06, range 0.24 to 0.68) involving 287 (79.7%) regions, and significant hypoconnectivity in 5,328 edges (8.3% of all edges; mean±SD Cohen’s *d*=−0.30±0.09, range −0.99 to −0.22) involving 356 (98.9%) regions after multiple comparisons at *P*FDR<0.05. That is, nearly all cortical brain regions were implicated in both forms of FC alterations in TLE patients. Hyperconnectivity and hypoconnectivity exhibited distinct spatial patterns. At the network level, hyperconnectivity mainly involved edges between the default mode and dorsal attention (*N*_norm_=19%; see **Methods** for definition of *N*_norm_), frontoparietal (*N*_norm_=12%), and ventral attention (*N*_norm_=10%) networks, and between limbic and dorsal attention networks (*N*_norm_=10%; **Figure 2b**). Conversely, hypoconnectivity was predominant between the default mode and limbic networks (*N*_norm_=26%), between the ventral attention and somatomotor networks (*N*_norm_=20%), and within the ventral attention network (*N*_norm_=23%; **Figure 2c**). At the regional level, hyperconnectivity clustered primarily in anterior lateral temporal, superior parietal, supramarginal gyrus, rostral middle frontal, and medial association cortices (superior frontal, orbitofrontal, and precuneus), which are key nodes of the default mode and dorsal attention networks (**Figure 2b**) involved in cognitive control and memory retrieval (**Supplementary Figure 2**). On the other hand, hypoconnectivity was concentrated primarily in medial (parahippocampus, entorhinal) and anterior lateral temporal (superior, middle, inferior), insular, caudal anterior/posterior cingulate, and paracentral cortices, which are temporolimbic and paralimbic territories (**Figure 2c**) implicated in episodic memory, attention, and arousal (**Supplementary Figure 2**).

**Figure 2.**
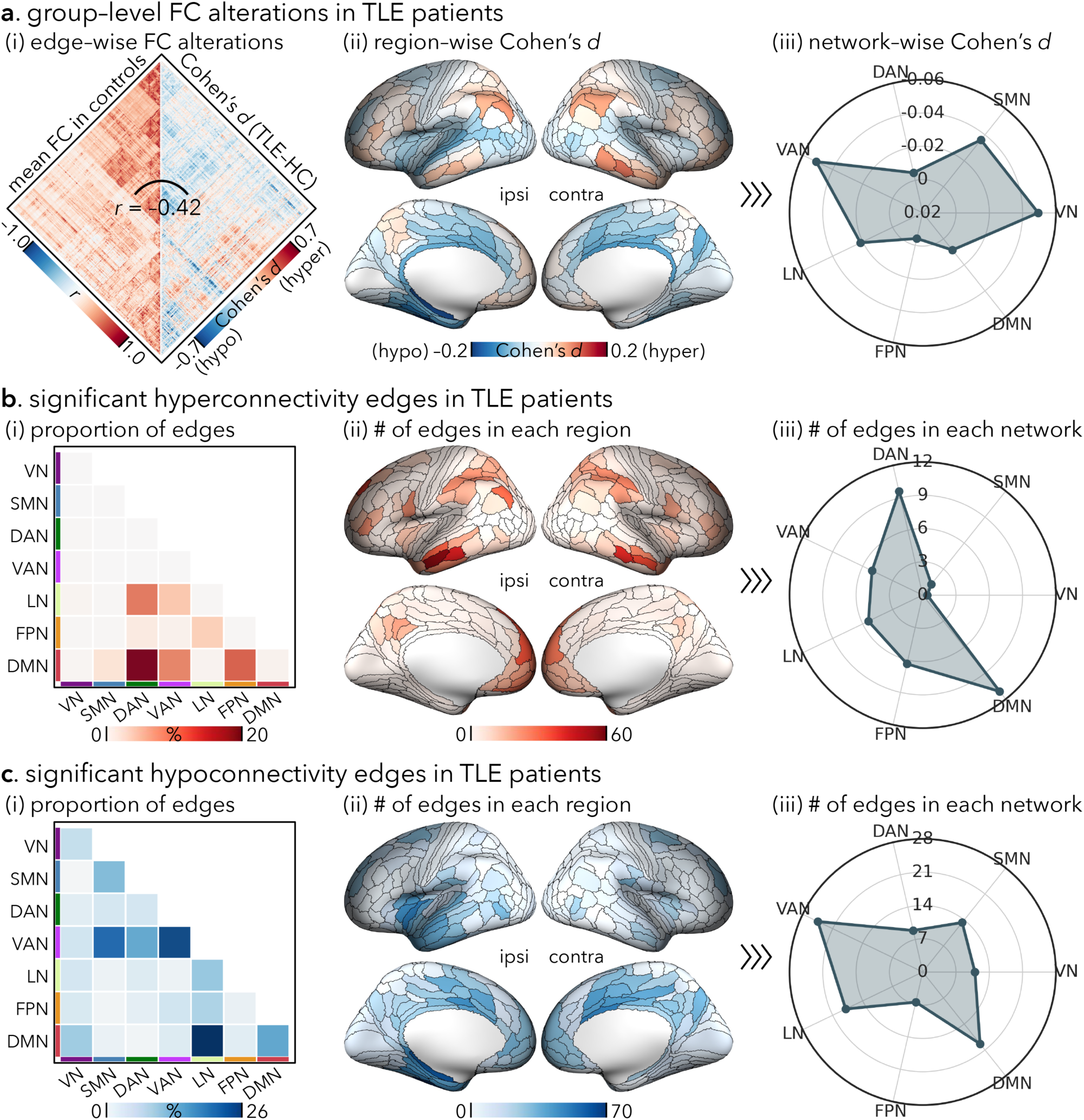
Group-level FC alterations in TLE patients. **(a)** FC differences between TLE patients (*n*=297) and healthy controls (*n*=282). **(i) Left**: Group-averaged FC matrix (360×360) in healthy controls. **Right**: Edge-wise group differences expressed as Cohen’s *d* (warm, hyperconnectivity; cool, hypoconnectivity). **(ii)** Region-wise mean Cohen’s *d*, averaged across all incident edges and projected onto the cortical surface (ipsi/contra indicate the hemisphere relative to the seizure focus). **(iii)** Network-wise mean Cohen’s *d*, averaged across brain regions within each of the seven canonical functional networks. **(b-c)** Significant **(b)** hyperconnectivity and **(c)** hypoconnectivity edges after false discovery rate (FDR) correction at a *P*FDR<0.05. **(i)** Proportion of significant edges within and between networks, normalized by the total number of possible edges in each network (i.e., *N*_norm_). **(ii)** Region-level degree of altered connectivity, quantified as the number of significant edges incident on each region. **(iii)** Network-level summary of altered connectivity, quantified as the average number of significant edges within each network. **Abbreviations**: ipsi, ipsilateral; contra, contralateral; VN, visual network; SMN, somatomotor network; DAN, dorsal attention network; VAN, ventral attention network; LN, limbic network; FPN, frontoparietal network; DMN, default mode network.

Findings were largely consistent across datasets. Within-dataset TLE–control analyses yielded effect-size maps (Cohen’s *d*) that correlated strongly with the full-sample discovery analysis (*r*=0.34–0.89, *P*<0.001) and across datasets (*r*=0.08–0.35, all *P*<0.001). Despite variability in the implicated edges, the overall patterns of hyperconnectivity and hypoconnectivity were reproducible across datasets and scanners (**Supplementary Figure 3**).

#### 2.2.2 Individual-Level FC Alterations

Patient-specific extreme FC deviations were identified by thresholding *z*-scored FC values relative to healthy controls. Specifically, extreme positive deviations (hyperconnectivity) were defined as edges with *z*>1.5, while extreme negative deviations (hypoconnectivity) were defined as edges with *z*<−1.5 (**Figure 3a**). These reflect the deviations from the healthy control distributions by at least 1.5 standard deviations at the individual level. We then quantified, for each edge, the number of patients showing extreme deviations. Across all edges, the median proportion of TLE patients exhibiting hyperconnectivity was 7.1% (IQR 5.1%–8.8%, range 0.1%–24.6%), whereas hypoconnectivity was observed in a median of 8.1% of patients (IQR 6.4%–10.4%, range 0.3%–41.8%). Thus, for a typical edge, ∼7.1% and ∼8.1% of individuals were considered hyperconnected and hypoconnected outliers, respectively. At the individual level, the counts of hyperconnectivity and hypoconnectivity edges were negatively correlated across patients (*r*=−0.33, *P*<0.001), reflecting substantial inter-individual variability, with some patients showing predominantly enhanced FC and others predominantly weakened FC.

**Figure 3.**
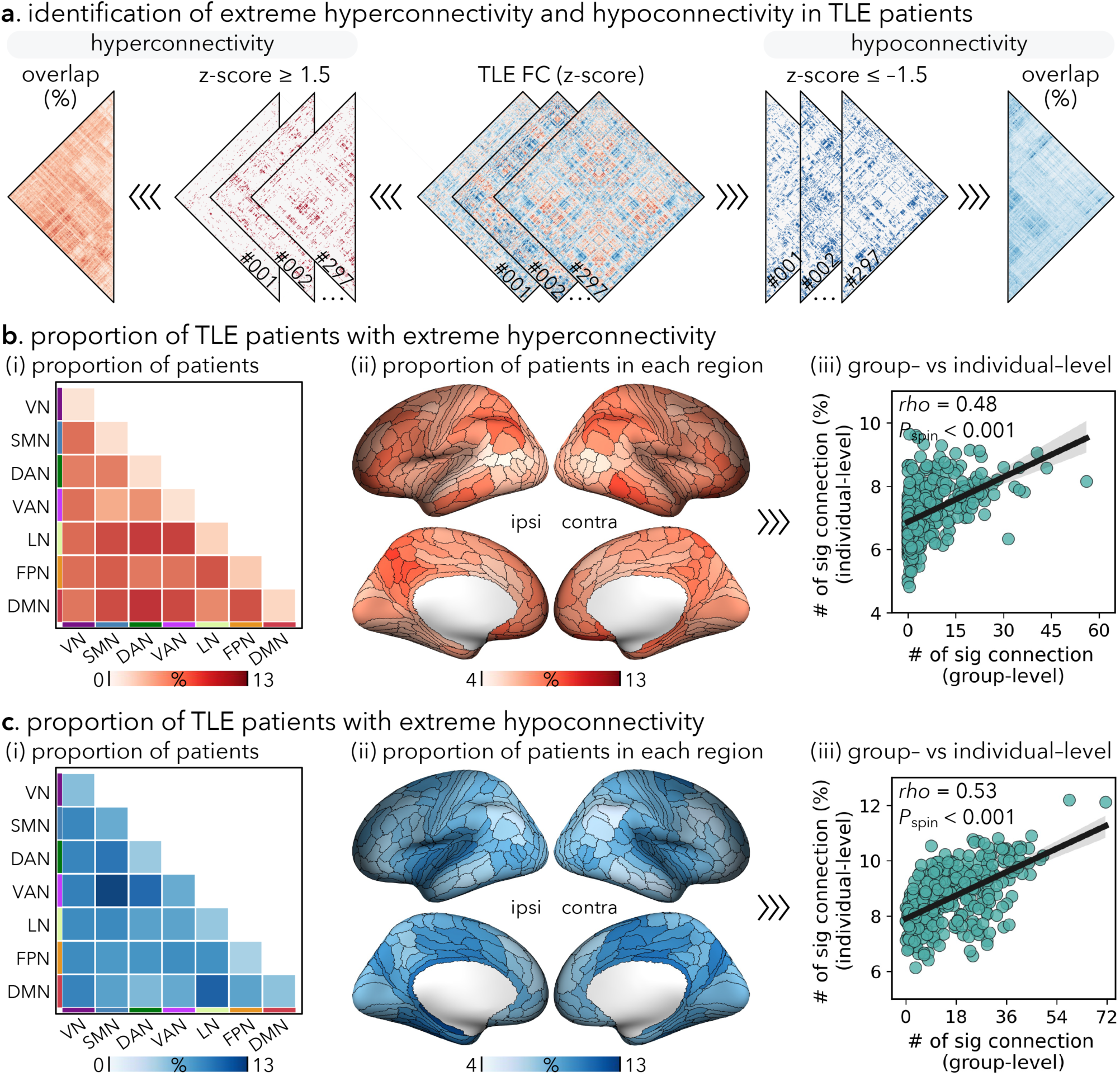
Individual-level FC alterations in TLE patients. **(a)** Schematic illustration of identifying extreme FC deviations in individual patients. Edge-wise FC *z*-scores (relative to healthy controls) were thresholded at |*z*|≥1.5 to identify hyperconnected (*z*≥1.5) and hypoconnected (*z*≤−1.5) edges. Binary outlier maps were then aggregated across patients to generate edge-wise overlap maps showing the proportion of patients exhibiting extreme deviations. **(b-c)** Spatial distribution of extreme **(b)** hyperconnectivity and **(c)** hypoconnectivity across patients. **(i)** Proportion of patients showing extreme deviations within and between networks. **(ii)** Regional burden of deviations, expressed as the proportion of patients showing extreme deviations in each brain region (ipsi/contra indicate hemisphere relative to the seizure focus). **(iii)** Correspondence between group-level and individual-level FC alterations across brain regions, relating the number of significant connections identified at the group level (x-axis; from Figure 2b-c) to the proportion of patients showing extreme deviations at the individual level (y-axis). Each dot represents one brain region; the solid line denotes Spearman’s rank correlation with a 95% CI (shaded band). Significance (*P*_spin_) was assessed with spin permutation tests (5,000 rotations; one-sided). **Abbreviations**: ipsi, ipsilateral; contra, contralateral; VN, visual network; SMN, somatomotor network; DAN, dorsal attention network; VAN, ventral attention network; LN, limbic network; FPN, frontoparietal network; DMN, default mode network.

At the network level, hyperconnectivity was most prevalent in edges between the default mode/limbic network and other networks (**Figure 3b**). In contrast, hypoconnectivity was most common in edges between the default mode/ventral attention network and other networks (**Figure 3c**). At the regional level, hyperconnectivity predominated in bilateral lateral and medial association cortices (**Figure 3b**), whereas hypoconnectivity was most pronounced in medial temporal, insular and cingulate cortices (**Figure 3c**). The spatial distributions of hyperconnectivity (*rho*=0.48, *P*_spin_<0.001) and hypoconnectivity prevalence (*rho*=0.53, *P*_spin_<0.001) mirrored the patterns observed in the group analysis (**Figure 3b-c**), indicating that the networks and regions exhibiting the strongest cohort-level abnormalities are also those most frequently implicated in individual patients.

### 2.3 Structural Determinants of FC Alterations in TLE

Building on the identified spatially heterogeneous hyperconnectivity and hypoconnectivity patterns, we next asked to what extent they are constrained by the brain’s multiscale architecture. We therefore derived three complementary inter-regional affinity metrics in healthy controls, previously commonly used to capture distinct components of cortical wiring collectively (*40*, *41*, *48*): (i) geodesic distance (GD), indexing interregional proximity along the cortical sheet (*49*); (ii) structural connectivity (SC), reflecting white matter streamlines between regions (*50*); and (iii) microstructural profile covariance (MPC), quantifying interregional similarity of intracortical microstructural profiles across cortical depths (**Figure 4a**) (*51*). For each brain region, we fitted a multilinear regression model that predicts the region’s cortex-wide FC alteration profile (i.e., Cohen’s *d*) from a linear combination of its structural affinity profiles (i.e., GD, SC and MPC). We found marked regional heterogeneity in model fits across the neocortex (adjusted *R*^2^=0.01–0.35), with the strongest fits in temporo-insular and lateral frontoparietal cortices, corresponding at the network level to the limbic (*R*^2^=0.13), frontoparietal (*R*^2^=0.12), and default mode (*R*^2^=0.11) networks. Dominance analysis, used to quantify the unique contribution of each structural affinity measure, revealed regionally variable dominance patterns, with the largest overall contributions at MPC (mean 46.6%, IQR 20.4%–73.3%), followed by GD (mean 31.8%, IQR 10.1%–50.7%) and SC (mean 21.6%, IQR 7.3%–30.0%; **Figure 4b**).

**Figure 4.**
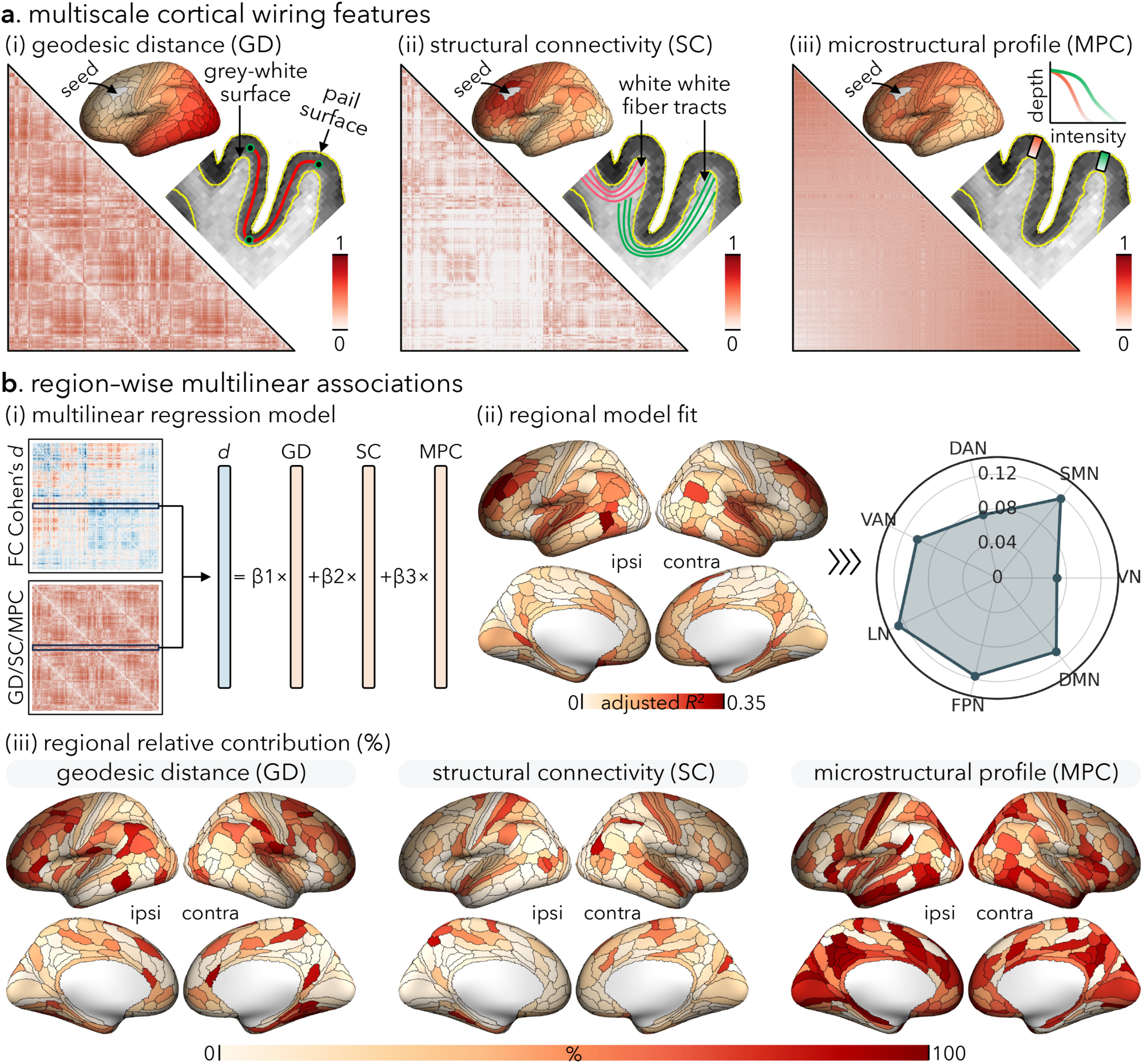
Structural determinants of FC alterations in TLE patients. **(a)** Multiscale cortico-cortical wiring features derived in healthy controls, including **(i)** geodesic distance (GD), **(ii)** structural connectivity (SC), and **(iii)** microstructural profile covariance (MPC). **(b)** Region-wise associations between structural affinity profiles and TLE-related FC alteration effect sizes using a multilinear regression model. **(i)** A multilinear regression model was applied to predict a region’s FC alteration profile (Cohen’s *d*) from its structural affinity profiles (GD, SC and MPC). **(ii) Left**: The distribution of model fits (adjusted *R*^2^) was projected onto the cortical surface (ipsi/contra indicate the hemisphere relative to the seizure focus). **Right**: Network-wise mean *R*^2^, averaged across brain regions within each of the seven canonical functional networks. **(iii)** The relative contribution (%) of each structural affinity profile across the cortex derived from dominance analyses. **Abbreviations**: ipsi, ipsilateral; contra, contralateral; VN, visual network; SMN, somatomotor network; DAN, dorsal attention network; VAN, ventral attention network; LN, limbic network; FPN, frontoparietal network; DMN, default mode network.

*Post hoc* analyses by relating these structural affinity profiles to FC alteration profiles using univariate Spearman’s rank correlations revealed predominantly positive associations across the cortex. That is, brain regions with stronger anatomical proximity, higher SC strength or greater microstructural similarity tended to exhibit more pronounced FC alterations in TLE patients (**Supplementary Figure 4**). Moreover, zooming in on individual regions, we also revealed the strongest correlations in the lateral prefrontal, temporal and insular cortices, which encompass key hubs of the default mode, frontoparietal, and paralimbic networks. Altogether, these results suggest that FC alterations in TLE preferentially align with the brain’s microscopic and macroscopic architecture specifically in the higher-order association cortices.

### 2.4 Clinical Relevance of FC Alterations in TLE

#### 2.4.1 Clinical Features

To identify FC patterns associated with disease severity, we applied partial least squares (PLS) analysis to relate patient-level FC measures on edges showing significant group-level changes to a set of clinical features. PLS identified one statistically significant latent variable (LV1), driven by hypoconnectivity (brain–clinical score correlation: *r*=0.19, *P*=0.001, *P*_perm_=0.017). Clinical loadings (i.e., correlations between individual clinical score and LV1) indicated that longer disease duration (*r*=−0.90), older age (*r*=−0.66), and the presence of antiseizure medication resistance (*r*=−0.41) and hippocampal sclerosis (*r*=−0.39) were associated with greater hypoconnectivity (**Figure 5a**).

**Figure 5.**
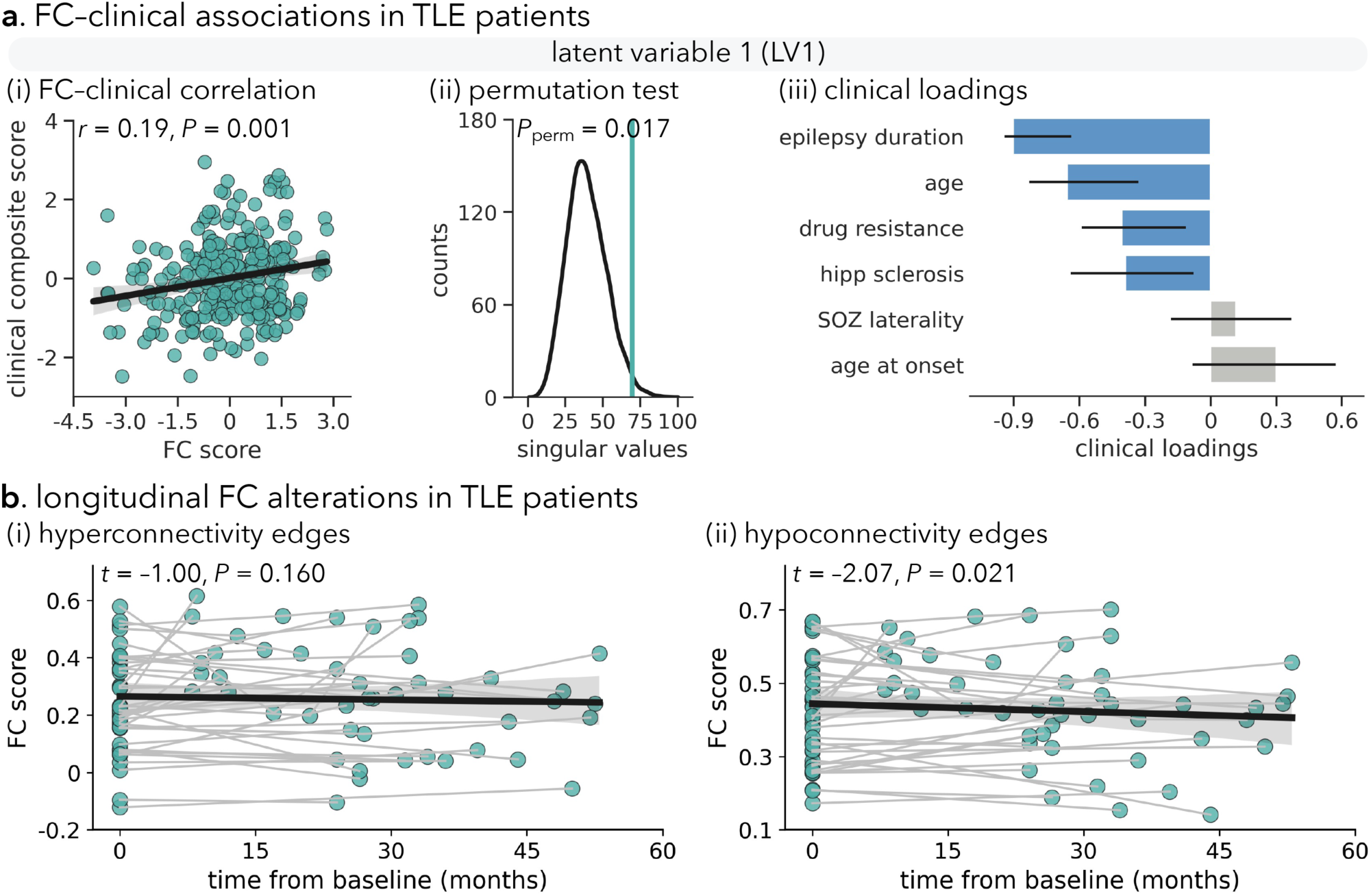
Clinical relevance and longitudinal trajectories of FC alterations in TLE. **(a)** Partial least squares (PLS) correlation analysis relating patient-level FC alterations to clinical features. **(i)** Association between FC scores and clinical composite scores for the first latent variable (LV1). Each point represents one patient; the solid line shows the Pearson correlation with a 95% confidence interval (shaded band). **(ii)** Permutation test for LV1, showing the null distribution of LV1 singular values obtained by shuffling clinical data 5,000 times; the vertical line denotes the observed singular value. **(iii)** Clinical loadings on LV1 with bars showing correlations between each clinical variable and LV1 scores. Error bars indicate 95% confidence intervals across 5,000 bootstrap resamples. Variables with reliable contributions are highlighted in blue. **(b)** Longitudinal FC trajectories in patients with repeated MRI scans. Patient-specific mean FC values on edges showing **(i)** hyperconnectivity and **(ii)** hypoconnectivity are plotted against time from baseline (months). Each dot represents one scan; lines connect repeated scans from the same individual, and the thick line indicates the mixed-effects trend with a 95% CI (shaded band). **Abbreviations**: hipp, hippocampus; SOZ, seizure onset zone.

#### 2.4.2 Longitudinal FC Alterations

We examined longitudinal alterations in FC among patients with ≥ 2 presurgical MRI scans (2 visits: *n*=39; 3 visits: *n*=8; 86 scans total; first-to-last scan interval: mean 29 months, range 8–53 months). Hypoconnected edges exhibited a further decrease in FC over time (*t*=−2.07, *P*=0.021; **Figure 5b**). Although hyperconnected edges demonstrated a trend toward lower FC, the effect was not significant (*t*=−1.00, *P*=0.160). Therefore, these within-patient trajectories demonstrate progressive network disruption in patients and support early identification of individuals transitioning toward a more disconnected network state.

#### 2.4.3 Postsurgical Seizure Outcome

Finally, we investigated whether presurgical FC alterations predict postsurgical seizure outcomes in 98 TLE patients who underwent unilateral temporal lobectomy. For each patient, extreme FC deviations were identified by thresholding the *z*-scored FC matrix at |*z*|≥1.5 (**Figure 6a**). Compared with patients rendered seizure-free (SF; *n*=73), patients with persistent postsurgical seizures (non-seizure-free, NSF; *n*=25) exhibited a greater number of hypoconnected edges (*t*=2.36, *P*FDR=0.022). In contrast, SF and NSF subgroups did not differ in seizure-focus laterality (*χ*²=0.33, *P*=0.568) or the presence of hippocampal sclerosis (*χ*²=1.12, *P*=0.289). Meaning that the FC difference was not explained by these clinical characteristics. Furthermore, to localize these effects, patient-specific surgical cavity mask was derived from postsurgical T1-weighted MRI, and whole-brain edges were categorized into resected–resected, resected–spared, and spared–spared sets. A significant difference between SF and NSF patients emerged only in spared–spared edges—that is, among “healthy” brain regions (*t*=−2.32, *P*FDR=0.012; **Figure 6b**). That is, greater FC reduction beyond the primary epileptogenic focus was associated with unfavourable postsurgical outcomes.

**Figure 6.**
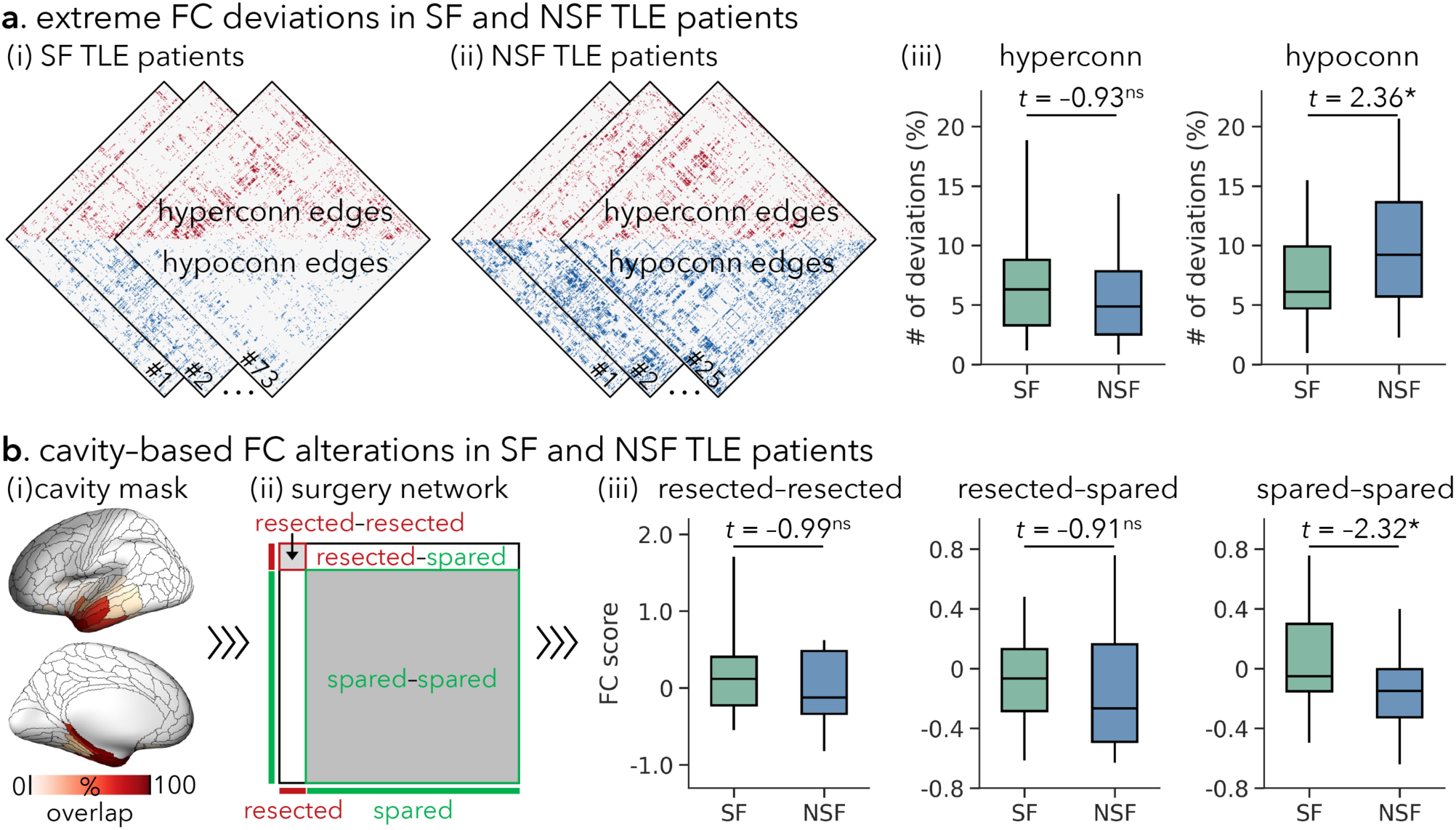
Presurgical FC alterations and postsurgical seizure outcome in TLE. **(a)** Extreme FC deviations stratified by postsurgical seizure freedom. **(i-ii)** Patient-wise extreme hyperconnectivity and hypoconnectivity edges for **(i)** seizure-free (SF) and **(ii)** non-seizure-free (NSF) patients, identified by thresholding each patient’s *z*-scored FC matrix at |*z*|≥1.5. **(iii)** Group comparisons between SF and NSF patients of the number of extreme hyperconnectivity and hypoconnectivity edges per individual. **(b)** Cavity-based FC alterations stratified by postsurgical seizure outcome. **(i)** Spatial overlap of the surgical cavity masks across TLE patients. **(ii)** Definition of surgery-related FC categories based on resected and spared regions: resected-resected, resected-spared, and spared-spared connections. **(iii)** Comparisons of individual mean presurgical FC values between SF and NSF patients within each category. Boxplots show the median line, interquartile range (25th−75th percentiles), and whiskers extending to 1.5× interquartile range. Asterisks denote significance at *P*<0.05 from two-sample *t*-tests. **Abbreviations**: hyperconn, hyperconnectivity; hypoconn, hypoconnectivity; ns, non-significant.

### 2.5 Disease Specificity of FC Alterations

Having identified robust, clinically relevant FC alterations in TLE, we next asked whether these patterns were specific to TLE or reflected general effects across focal epilepsy. To address this question, we compared TLE with an independent cohort of extra-TLE focal epilepsy. Although both TLE and extra-TLE cohorts showed extensive network-level FC alterations relative to healthy controls, extra-TLE patients showed greater FC changes of the visual (mean Cohen’s *d*=−0.12) and attentional (mean Cohen’s *d*=−0.15) networks. Similarly, region-level analyses showed that FC changes in extra-TLE patients were more pronounced in the precentral, inferior parietal, and occipital cortices. Direct comparisons between TLE and extra-TLE patients further highlighted these dissociations, revealing distinct spatially organized FC profiles across epilepsy syndromes (**Supplementary Figure 5**). Therefore, these FC alterations identified in TLE are not solely attributable to epilepsy *per se* but instead reflect syndrome-specific patterns of large-scale functional network reorganization.

## 3. Discussion

This study represents one of the largest multicentre pooled analyses to date of intrinsic FC alterations in TLE. Using whole-brain FC analysis, we identified a robust, hub-centred pattern of network reorganization, characterized by hyperconnectivity in frontoparietal association cortices and hypoconnectivity in medial/lateral temporal and paralimbic regions. Crucially, these alterations were not spatially random, but markedly constrained by the cortex’s structural scaffold, including corticocortical proximity, microstructural similarity, and white matter connectivity. Furthermore, the FC phenotype was shown to be closely associated with disease burden and postsurgical seizure outcome, and reliably distinguished TLE from other focal epilepsies. These results collectively demonstrate a robust signature of intrinsic functional network disruption in TLE that is partially shaped by the cortical structural wiring and has potential utility for whole-brain pathophysiological profiling and for informing pre-surgical evaluation and outcome prognostication for individual patients.

By aggregating multiple independent cohort datasets, the present study benefited from high statistical power, enabling the identification of robust and reproducible intrinsic FC changes in TLE. We found that regions and networks of dysfunction consistently converged on high-order association systems, including the default mode, frontoparietal, dorsal and ventral attention, and limbic networks. Notably, nearly all regions implicated in our findings have previously been characterized as rich-club nodes (densely interconnected hubs), particularly the medial prefrontal, precuneus, anterior and posterior cingulate, lateral temporoparietal, and insular regions (*52-54*). Hub regions are considered core nodes in brain networks and promote global communication and integrative information processing within and across diverse systems (*55*). On the other hand, dense connections and topological centrality render hubs particularly susceptible to pathological disturbances, such as network overload and hyper-synchronization (*56-58*). In the context of TLE, such vulnerability can facilitate maladaptive reorganization of distributed circuits, leading to extensive FC changes. In alignment with our findings, prior studies using high-resolution structural MRI have shown that grey matter loss and structural network reorganization associated with TLE are not confined to the mesial temporal lobe but instead preferentially affect hubs across the neocortex (*59-62*). Importantly, our hub-centred dysfunction pattern was highly consistent across datasets and patients with differing acquisition contexts and clinical compositions, underscoring its crucial role as a syndrome-general network substrate in TLE rather than as idiosyncratic effects driven by individual disease histories. Together, convergent evidence from group and individual analyses provides strong support for the role of temporolimbic and adjacent transmodal association cortices in TLE pathophysiology (*44*, *63-65*). This broadly extends prior rs-fMRI studies that have often been constrained to regions or connections of interest due to limited sample sizes and statistical power, by offering a connectome-wide, reproducible profile of functional network vulnerability in TLE.

Multimodal MRI acquisitions, such as those aggregated in the present study, enable integrated characterization of multiple *in vivo* features of cortical wiring, allowing us to investigate how intrinsic FC alterations in TLE are shaped by multiscale structural architecture. By combining geodesic distance, white matter tractography strength, and microstructural profile covariance, we captured complementary aspects of cortical connectivity that converge on a common principle of communication capacity: regions that are closer in physical distance, have more direct fibre pathways, have more similar intra-cortical microstructures, and are inherently predisposed to interact and maintain coordinated dynamics (*40*, *41*). In fact, geodesic distance along the cortical surface is considered to reflect the cost of short-range horizontal connectivity; spatially proximal regions can establish interactions at lower wiring and metabolic cost (*49*, *66*). Tract length indexes short- and long-range white matter fibres (*50*). Microstructural profile covariance reflects similarity in intracortical laminar differentiation and cyto-architectural organization, which is linked to hierarchical localization and intrinsic computational properties (*51*, *67*). In TLE, high structural constraints on altered FC observed in the prefrontal and temporal cortices are likely due to their distinctive roles in cortical organization. These regions are commonly subsumed under the umbrella of transmodal association cortex, characterized by less pronounced myelination, prolonged maturation, and longer-range connectivity patterns, across large-scale systems (*68-71*). At the top of the cortical hierarchy, frontotemporal association cortices act as integrative hubs that coordinate information across systems (*53*, *72*, *73*). Therefore, their intrinsic functional architecture is tightly anchored to the underlying structural scaffold, making functional reconfiguration most likely to unfold along these pre-existing communication backbones (*35*, *36*, *39*, *74*). From a systems-level perspective, this organization renders the frontotemporal association networks particularly vulnerable to distributed functional reconfigurations in TLE, such that intrinsic functional changes are not expressed uniformly across the cortex but instead preferentially emerge where dense structural embedding amplifies network-level perturbations. Together, our findings support a model in which large-scale functional network reorganization in TLE is guided by intrinsic wiring principles of the cortex, linking multiscale structural organization to system-level functional alterations, with the strongest constraints expressed in the frontotemporal association regions.

Clinical analyses further revealed convergent associations between intrinsic FC alterations and disease duration, progression, and post-surgical seizure outcomes, particularly in the medial and lateral temporal and paralimbic cortices. Specifically, patients with long-standing epilepsy showed greater FC decreases, reflecting progressive degradation of large-scale functional coupling over the course of TLE (*24*, *75*). This pattern aligns with prior cross-sectional morphometric and tractography assessments, which show that multilobar grey and white matter changes associated with TLE increase with longer disease duration (*32*, *60*, *76*, *77*). Thus, the observed functional changes may have long-lasting consequences for large-scale network integration, reinforcing the case for early identification and surgical intervention in pharmacoresistant TLE, as earlier therapy may improve seizure outcomes and clinical efficacy (*78-80*). We also observed more pronounced FC declines in temporolimbic cortices among patients with pharmacoresistant TLE than in those with better-controlled seizures. This suggests that intrinsic functional alterations may serve as a valuable tool for predicting the effectiveness of antiseizure drugs, which remains challenging to date (*81*, *82*). Nevertheless, the observation warrants cautious interpretation, as influential treatment-related confounds, including dosage, duration, and the number of previous antiseizure drug trials, were not available and were not considered in the study. Future studies incorporating more comprehensive treatment histories will be essential to provide a more nuanced delineation.

Finally, our findings indicate that large-scale functional network alterations on presurgical imaging may be relevant to postsurgical outcomes in TLE. A direct comparison between SF and NSF TLE subcohorts revealed greater FC decreases in the NSF TLE subcohort. Strikingly, *post hoc* evaluation across the entire cortex showed a specific association between persistent postsurgical seizures and FC decline in regions outside the mesiotemporal lobe. This observation aligns with recent structural network studies on grey-matter morphology, which similarly reported that SF and NSF patients differed primarily in neocortical regions remote from the presumed epileptogenic zone (*83*). These convergent findings on function and structure echo the idea that an unfavourable surgical outcome is driven by pathology beyond the seizure focus (*23*, *84*). This network-level mechanism provides a plausible explanation for the persistent disabling seizures postsurgically in 30%–50% of TLE patients. Traditionally, this phenomenon has been attributed mainly to inaccurate or incomplete localization and lateralization of seizure onset zones before surgery. Our results instead emphasize that mapping the configurations of epileptogenic networks, together with residual networks, is crucial for reliably determining seizure outcomes. Thus, functional MRI based on connectomes may complement conventional structural MRI markers, particularly hippocampal atrophy, and substantially enhance current prediction models. Significantly, identifying patients with network configurations associated with a high likelihood of seizure freedom after unilateral resection can facilitate more confident surgical decision-making and potentially reduce the need for invasive intracranial monitoring. Collectively, these findings underscore the clinical significance of mapping distributed functional network alterations to improve outcome stratification and optimize individualized surgical planning in focal epilepsy.

## 4. Methods

### 4.1 Participants

This multicentre, retrospective study included 652 participants aged 18–70 years: 297 patients with unilateral TLE (137 [46.1%] males; mean±SD age=32.8±10.5 years, range 18–68.2 years; 153 [51.5%] left TLE), 73 disease controls with unilateral extra-TLE focal epilepsy (36 [49.3%] males; mean±SD age=30.3±11.8 years, range 18–69.3 years; 23 [50.7%] left extra-TLE), and 282 age- and sex-matched healthy controls (142 [50.1%] males; mean±SD age=29.3±8.1 years, range 18–60 years). Participants were aggregated from four independent datasets at three epilepsy centres: (i) Montreal Neurological Institute–Hospital (MICA-MICs: 72 TLE, 45 extra-TLE, 100 healthy controls; NOEL: 72 TLE, 28 extra-TLE, 42 healthy controls) (*45*), (ii) Universidad Nacional Autónoma de México (EpiC: 29 TLE, 34 healthy controls) (*46*), and (iii) Jinling Hospital, Nanjing University School of Medicine (Nanj: 124 TLE, 106 healthy controls) (*23*). This study was approved by the research Ethics Committees of each centre (MICA-MICs and NOEL: Montreal Neurological Institute–Hospital, McGill University; EpiC: Institute of Neurobiology, Universidad Nacional Autónoma de México; Nanj: Jinling Hospital, Nanjing University School of Medicine). All participants provided written informed consent in accordance with the Declaration of Helsinki.

Epilepsy diagnosis and lateralization were established by epilepsy specialists at each centre using the International League Against Epilepsy (ILAE) seizure and syndrome criteria. Each patient underwent a comprehensive presurgical multidisciplinary evaluation—including clinical history, ictal semiology, prolonged video-EEG telemetry (ictal and interictal), and clinical MRIs (and PET when available). Exclusion criteria included mass lesions (tumours, malformations of cortical development, vascular malformations), traumatic brain injury, encephalitis, and comorbid neurological or psychiatric disorders. Clinically, 76.8% (*n*=225) of TLE patients and 86.1% (*n*=62) of extra-TLE patients had persistent seizures. TLE patients had a mean age at seizure onset of 18.1 years (SD 10.8 years, range 0.3–60 years) and a mean epilepsy duration of 14.4 years (SD 11.0 years, range 0.1–49 years). Extra-TLE patients had a mean age at seizure onset of 13.0 years (SD 9.9 years, range 0.5–66 years) and a mean epilepsy duration of 16.5 years (SD 12.5 years, range 0.5–65.3 years). Among TLE patients, 66.3% (*n*=197) had ipsilateral hippocampal sclerosis, as confirmed by radiological reports, quantitative analyses of T1-weighted and FLAIR images (hippocampal volume loss and/or hyperintensity), and, when available, postsurgical histopathological examination (*85-87*); 8.8% (*n*=26) had ipsilateral hippocampal gliosis or focal cortical dysplasia; 24.9% (*n*=74) were MRI-negative. At the time of analysis, 98 (33.0%) TLE patients had undergone unilateral temporal lobectomy. Postsurgical seizure outcome was assessed using the Engel classification system (*88*): 73 (74.5%) were seizure-free (Engel I) and 25 (25.5%) had persistent postsurgical seizures (Engel II–IV), with an average follow-up of 44 months (SD 34 months, range 12–108 months). Demographic and clinical information for participants in each dataset is provided in **Table 1**.

### 4.2 MRI Acquisition and Processing

Multimodal MRI data, including T1-weighted, rs-fMRI, and diffusion MRI, were acquired on 3-Tesla scanners for all participants (before surgery for patients; MICA-MICs: Siemens Prisma; EpiC: Philips Achieva; NOEL and Nanj: Siemens Trio). Quantitative T1 (qT1) relaxometry data were also available in the MICA-MICs cohort. 39 TLE patients (MICA-MICs: *n*=22; EpiC: *n*=17) had repeated scans (2 visits: *n*=39; 3 visits: *n*=8) on the same scanners, with a mean interval between the first-to-last scans of 29 months (SD 14 months, range 8–53 months); 86 serial scans were available for the longitudinal analysis. Postsurgical T1-weighted images were available from 52 TLE patients with unilateral temporal lobectomy. Multimodal MRI data for the four datasets were uniformly processed with *micapipe* (v0.2.3; http://micapipe.readthedocs.io), an openly accessible multimodal MRI processing and fusion pipeline (*89*). Briefly, for rs-fMRI data, the first five volumes were excluded to ensure steady-state magnetization. Six head motion parameters, spike regressors, and signals of white matter and cerebrospinal fluid were regressed out. The whole cortex was parcellated into 360 cortical brain regions using the hemisphere-matched HCP-MMP1.0 atlas (*47*), and surface-based time series were averaged across all vertices within each region. Details of MRI acquisition and processing are provided in the **Supplementary Materials**.

### 4.3 FC Alteration Analysis in TLE

#### 4.3.1 Group-Level FC Alterations

For each participant, Pearson’s correlation coefficients were computed between time series across all regions, yielding a 360×360 FC matrix with 64,620 unique edges. To mitigate site-related batch effects, edge-wise FC values were harmonized across datasets using ComBat with empirical Bayes (*90*), while preserving effects of age, sex, and diagnosis. Each patient’s FC matrix was normalized relative to the healthy control group to generate a 360×360 *z*-scored matrix, quantifying deviations from the control mean (positive/negative *z*-scores indicating increased/decreased FC). For right TLE, FC matrices were flipped left to right to align seizure laterality across patients (ipsilateral versus contralateral). TLE–control FC difference at the edge level was assessed using general linear models, including age and sex as covariates, with effect sizes expressed as Cohen’s *d*. Findings were corrected for multiple comparisons using the false discovery rate (FDR) with a *P*FDR<0.05. Significantly altered edges were characterized at three spatial scales: (i) the edge level; (ii) the network level, aggregating regions into seven canonical functional networks defined in the Yeo atlas (**Supplementary Figure 6**) (*10*); and (iii) the region level, quantifying the number of significantly altered edges linked to each brain region. To account for differences in network size, network-level characterization was expressed as the proportion of affected edges within each network, normalized by the total number of possible edges in that network (i.e., *N*_norm_). Cognitive and behavioral terms related to FC alterations were decoded using the Neurosynth (https://neurosynth.org/) (*91*) by computing the spatial similarity between hyper/hypoconnectivity maps (**Figure 2b-c**) and probabilistic activation maps for cognitive/behavioral terms. Lastly, replicability of FC alterations across datasets was assessed by (i) repeating the connectome-wide FC analysis and (ii) comparing subject-specific FC values of significantly altered edges, independently within each dataset (**Supplementary Figure 3**).

#### 4.3.2 Individual-Level FC Alterations

For each patient, edge-wise FC *z*-scores (relative to healthy controls) were thresholded at |*z*|≥1.5 to identify extreme hyperconnectivity (*z*≥1.5) and hypoconnectivity (*z*≤−1.5) deviations (**Figure 3a**). We then computed, for each edge, the proportion of patients exhibiting deviations. Results were summarized at the network-level as the mean proportion of deviations per network and at the region-level as the mean proportion of deviations per region. The correspondence between individual- and group-level FC alterations across regions was evaluated using Spearman’s rank correlation. The significance of spatial correlation was assessed using nonparametric spin-permutation tests (*92*). Specifically, parcels’ coordinates were randomly rotated on the sphere while preserving spatial autocorrelation, and values were reassigned to the nearest parcels. This procedure was repeated 5,000 times to generate null distributions. The *P*-value (*P*spin) was defined as the proportion of null correlations equal to or exceeding the empirical correlation (i.e., greater for positive correlations or weaker for negative correlations).

### 4.4 Associations with Cortical Structural Topography

To examine how FC alterations in TLE are spatially constrained by the brain’s structural architecture, we computed three complimentary features of cortical wiring in healthy controls: GD, SC, and MPC (**Figure 4a**) (*40*, *41*). GD, based on T1-weighted MRI and calculated as the shortest path between regional centroids along the cortical midsurface, reflects the spatial proximity and cortical wiring costs of two regions (*49*). SC, based on tractography applied to diffusion MRI, is an estimate of the white matter fiber bundles between each pair of regions. MPC, calculated as the inter-regional similarity of intracortical microstructural profiles sampled along 14 cortical columns from quantitative MRI, reflects laminar differentiation and cytoarchitectural complexity (*51*). Details of matrix construction are provided in the **Supplementary Materials**.

A multilinear regression model was applied for each brain region (**Figure 4b**), predicting the region’s FC alteration profile from a linear combination of its structural network profiles (GD, SC and MPC):

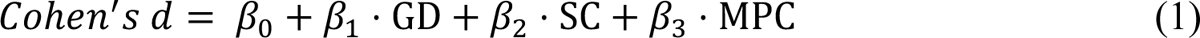

where the dependent variable was a row of the group-level FC difference matrix (i.e., Cohen’s *d*) and the independent variables (predictors) were the corresponding rows of the GD, SC and MPC matrices. The intercept *β*_0_ and the regression coefficients (*β*_1_, *β*_2_, and *β*_3_) were optimized to maximize the spatial correlation between the empirical and predicted Cohen’s *d* vectors. Goodness of model fit was quantified using adjusted-*R*^2^ (coefficient of determination). The relative contributions of independent variables were determined using dominance analysis. Briefly, dominance analysis constructed all possible combinations of predictors and refitted the multilinear model for every combination. The relative contribution of each predictor was defined as the increase in explained variance (i.e., gain in adjusted-*R*^2^) by adding that predictor to the models. Dominance scores were normalized by the total model fit, enabling comparisons across models.

In addition, to quantify the direct associations between FC alterations and structural network profiles across the cortex, we correlated each region’s FC Cohen’s *d* vector with the corresponding vector for each structural descriptor using Spearman’s rank correlation, yielding one correlation coefficient per region per measure (**Supplementary Figure 4**).

### 4.5 Clinical Relevance of FC Alterations in TLE

#### 4.5.1 Clinical Features

Associations between FC alterations and clinical features were examined using multivariate partial least squares (PLS) analysis. Mean FC *z*-scores for each patient were computed across edges showing significant FC increases and decreases separately (see **Figure 2b-c**). These FC measures were related to a clinical feature set comprising age, age at seizure onset, epilepsy duration, hippocampal sclerosis, antiseizure medication resistance, and seizure focus laterality. PLS analysis identified latent variables that maximized the covariance between FC and clinical features, yielding paired subject scores whose correlation quantified FC-clinical associations. The significance of latent variables was assessed with permutation tests, in which clinical features were shuffled across patients (5,000 times). The permutation *P*-value (*P*_perm_) was defined as the proportion of permuted singular values equal to or exceeding the observed value. The contribution of each clinical feature was evaluated with bootstrap resampling (5,000 repetitions), with bootstrap ratios computed as each feature’s weight divided by its bootstrap standard error.

#### 4.5.2 Longitudinal FC Alterations

Longitudinal FC alterations were analyzed in 39 patients with repeated MRI scans (2 visits: *n*=39; 3 visits: *n*=8; 86 serial scans in total) using linear mixed-effects models. *Time* (months since the baseline scan) was modelled as a fixed effect, with a random intercept for *subjects*, separately applied to the subject-specific mean FC across edges showing hyperconnectivity and hypoconnectivity. We specifically tested for the negative effects of *time* on FC.

#### 4.5.3 Postsurgical Seizure Outcome

The patient-specific surgical cavity was manually segmented by registering presurgical and postsurgical T1-weighted MRIs to the MNI152 standard template using linear transformations and then subtracting the postsurgical scan from the presurgical scan. Segmented cavities were visually inspected, manually edited, and mapped onto the fsLR-32k surface template. Cortical brain regions were partitioned into “resected” and “spared” territories (**Figure 6b**). FC edges were grouped into three categories: (i) resected–resected FC (edges between resected regions), (ii) resected–spared FC (edges between resected and spared regions), and (iii) spared–spared FC (edges between spared regions). Individual mean presurgical FC values were computed for each category and compared between SF-TLE and NSF-TLE subgroups using two-sample *t*-tests and FDR correction.

### 4.6 Disease Specificity of FC Alterations

To determine whether the identified FC alterations were specific to TLE, we performed parallel analyses in an independent cohort of patients with unilateral extra-TLE focal epilepsy, serving as a disease control group. For each edge, we conducted (i) extra-TLE versus healthy controls and (ii) TLE versus extra-TLE contrasts, controlling for age and sex. To summarize syndrome-specific findings, we contextualized edge-wise effect sizes (Cohen’s *d*) at (i) the network level by aggregating regions into seven canonical functional networks and (ii) the regional level by averaging Cohen’s *d* across incident edges for each region. We then projected these values onto the cortical surface. For a direct syndrome comparison, we also derived a TLE versus extra-TLE contrast (Cohen’s *d*) and displayed corresponding network- and region-level summaries (**Supplementary Figure 5**).

## Acknowledgements

K.X. is funded by the China Scholarship Council and the Savoy Foundation. J.C. and E.S. are supported by the Canadian Institutes of Health Research (CIHR) Vanier Canada Graduate Scholarships. A.N. is funded by the CIHR. R.R.C. is funded by the Fonds de Recherche du Québec - Santé (FRQ-S). S.L. is funded by the Centre de Recherche du CHUS. L.C. is funded by the Consejo Nacional de Ciencia y Tecnología (CONACYT) (181508, 1782, FC218-2023) and Dirección General de Asuntos del Personal Académico (DGAPA) - UNAM (IB201712, IG200117, IN204720, IN213423). B.C.B. acknowledges research support from the National Science and Engineering Research Council of Canada (NSERC RGPIN-2025-05932), CIHR (FDN-154298, PJT-174995, PJT-191853), SickKids Foundation (NI17-039), Helmholtz International BigBrain Analytics and Learning Laboratory (HIBALL), Healthy Brains and Healthy Lives (HBHL), Brain Canada Foundation, FRQS, Tier-2 Canada Research Chairs Program, and The Centre for Excellence in Epilepsy at the Neuro (CEEN).

## Data Availability

Raw data for MICA-MICs are available on the Canadian Open Neuroscience Platform (https://portal.conp.ca/) and the Open Science Framework (https://osf.io/j532r/). Raw data for EpiC are available on OpenNeuro (dataset ds004469, https://openneuro.org/datasets/ds004469/versions/1.1.4).

## Code Availability

MRI preprocessing was conducted using *micapipe* (v0.2.3; http://micapipe.readthedocs.io). Code for spin permutation test is available at https://github.com/frantisekvasa/rotate_parcellation or ENIGMA Toolbox (v2.0.1; https://enigma-toolbox.readthedocs.io). Code for image-based decoding is available at NiMARE (https://nimare.readthedocs.io) or BrainStat (v0.5.1, https://brainstat.readthedocs.io).

## Author Contributions

Study Concept/Design: K.X. and B.C.B. Data Acquisition, Analysis, Interpretation: K.X., J.C., E.S., F.F., T.A., R.R.C., M.N., A.N., A.D., R.P., S.L., Z.Z., L.C., A.B., N.B., and B.C.B. Writing – Original Draft: K.X. and B.C.B. Writing – Review & Editing: K.X., J.C., E.S., F.F., T.A., R.R.C., M.N., A.N., A.D., Alex.B., S.A., R.P., S.L., A.H., A.G.W., S.O., R.D., D.V.S., Z.Z., L.C., A.B., N.B., and B.C.B. Resources: Z.Z., L.C., A.B., N.B., and B.C.B. Supervision: A.B., N.B., and B.C.B.

## Supplementary Material

### MRI Acquisition

#### MICA-MICs dataset

Data were collected on a 3-T Siemens Magnetom Prisma-Fit scanner equipped with a 64-channel head coil, and included: (i) two T1-weighted scans (3D-MPRAGE, repetition time [TR] = 2300ms, echo time [TE] = 3.14ms, flip angle [FA] = 9°, field of view [FOV] = 256×256mm^2^, voxel size = 0.8×0.8×0.8mm^3^, matrix size = 320×320, 224 slices), (ii) one rs-fMRI scan (multiband accelerated 2D-BOLD echo-planar imaging [EPI], TR = 600ms, TE = 30ms, FA = 52°, FOV = 240×240mm^2^, voxel size = 3×3×3mm^3^, multi-band factor = 6, 48 slices, 700 volumes), (iii) one multi-shell diffusion MRI scan (2D spin-echo EPI, TR = 3500ms, TE = 64.40ms, FA = 90°, FOV = 224×224 mm^2^, voxel size = 1.6×1.6×1.6mm^3^, 3 b0 images, b-values = 300/700/2000s/mm^2^ with 10/40/90 diffusion directions), and (iv) one quantitative T1 scan (3D-MP2RAGE, TR = 5000ms, TE = 2.9ms, FA = 4°, FOV = 256×256mm^2^, voxel size = 0.8×0.8×0.8mm^3^, 240 slices). Participants were instructed to remain still, look at a cross, and not to fall asleep during rs-fMRI acquisition.

#### NOEL dataset

Data were collected on a 3-T Siemens Trio scanner equipped with a 32-channel head coil, and included: (i) one T1-weighted scan (3D-MPRAGE, TR = 2300ms, TE = 2.98ms, FA = 9°, voxel size = 1×1×1mm^3^), (ii) one rs-fMRI scan (2D gradient-echo EPI, TR = 2020ms, TE = 30ms, FA = 90°, voxel size = 4×4×4mm^3^, 34 slices, 150 volumes), and (iii) one diffusion MRI scan (2D twice-refocused EPI, TR = 8400ms, TE = 90ms, FA = 90°, voxel size = 2×2×3mm^3^, 63 slices, 1 b0 images, b-value = 1000s/mm^2^, 64 diffusion directions). Participants were instructed to keep their eyes closed and not to fall asleep during rs-fMRI acquisition.

#### EpiC dataset

Data were collected on a 3-T Philips Achieva scanner equipped with a 64-channel head coil, and included: (i) one T1-weighted scan (3D spoiled gradient-echo, TR = 8.1ms, TE = 3.7ms, FA = 8°, FOV = 256×256mm^2^, voxel size = 1×1×1mm^3^, 240 slices), (ii) one rs-fMRI scan (gradient-echo EPI, TR = 2000ms, TE = 30ms, FA = 90°, voxel size = 2×2×3mm^3^, 34 slices, 200 volumes), and (iii) one diffusion MRI scan (2D EPI, TR = 11.86s, TE = 64.3ms, FOV = 256×256mm^2^, voxel size = 2×2×2mm^3^, 2 b0 images, b-value = 2000s/mm^2^, 60 diffusion directions). Participants were instructed to keep their eyes closed and not to fall asleep during rs-fMRI acquisition.

#### Nanj dataset

Data were collected on a 3-T Siemens Trio scanner equipped with a 32-channel head coil, and included: (i) one T1-weighted scan (3D-MPRAGE, TR = 2300ms, TE = 2.98ms, FA = 9°, FOV = 256×256mm^2^, voxel size = 0.5×0.5×1mm^3^), (ii) one rs-fMRI scan (2D gradient-echo EPI, TR = 2000ms, TE = 30ms, FA = 90°, FOV = 240×240mm^2^, voxel size = 3.75×3.75×4mm^3^, 30 slices, 255 volumes), and (iii) one diffusion MRI scan (2D spin-echo EPI, TR = 6100ms, TE = 93ms, FA = 90°, FOV = 240×240mm^2^, voxel size = 0.94×0.94×3mm^3^, 4 b0 images, b-value = 1000s/mm^2^, 120 diffusion directions). Participants were instructed to keep their eyes closed and not to fall asleep during rs-fMRI acquisition.

### MRI Preprocessing

Multimodal MRI data were preprocessed using *micapipe* (v0.2.3; https://micapipe.readthedocs.io/) (*1*), an openly accessible multimodal MRI preprocessing and fusion pipeline that integrates AFNI, ANTs, FreeSurfer, FSL, MRtrix, and Workbench (*2-6*).

T1-weighted MRI data were de-obliqued, reoriented to standard LPI orientation (left to right, posterior to anterior, and inferior to superior), linearly co-registered, corrected for intensity non-uniformity, intensity normalized, and skull stripped. Subject-specific white and pial surfaces were generated from each native T1-weighted MRI by tracing tissue boundaries with FreeSurfer. Segmentation errors were manually corrected by placing control points. Native surfaces were then registered to the hemisphere-matched fsLR-32k template surface (32k vertices per hemisphere). The geodesic distance (GD), defined as the shortest path between two regions along the cortical surface, was computed (*7*). A centroid vertex was first defined for each region by identifying the vertex with the shortest summed Euclidean distance from all other vertices within its assigned region. Then, GD between the centroid vertex and all other regions was computed along the midthickness surface using Dijkstra’s algorithm (*8*).

Resting-state fMRI preprocessing included discarding the first five volumes, reorientation, slice-timing correction, head motion and distortion correction, and temporal band-pass filtering (0.01–0.08 Hz). Motion correction was performed using a rigid-body model, registering each timepoint volume to the mean volume. Motion outlier volumes (timepoints with prominent motion-related spikes) were identified and discarded with FSL’s motion outlier detection (*fsl_motion_outliers*) (*5*), using the default cutoff (75th percentile + 1.5× interquartile range). To maintain temporal continuity of the time series, we filled these censored frames with linear interpolation. Nuisance signal removal was performed using an in-house trained ICA-FIX classifier (for MICA-MICs) (*9*) or through linear regression of white matter and cerebrospinal fluid signals for other datasets (NOEL, EpiC, and Nanj). FreeSurfer surfaces were nonlinearly co-registered to the native average volumetric time series using label-based registration. Cortical time series were mapped from native fMRI space to the fsLR-32k surface template using trilinear interpolation, followed by spatial smoothing along the cortical mid-thickness sheet with a Gaussian kernel with a full width at half maximum (FWHM) of 10 mm. This surface-constrained smoothing improves signal-to-noise ratio, decreases variability, and retains sensitivity while limiting partial-volume blurring across tissue boundaries (*10-12*). Lastly, vertex-wise time series were parcellated into 360 cortical brain regions by averaging vertices within each region using the HCP-MMP1.0 atlas (*13*), a multimodal cortical parcellation comprising 180 homologous regions per hemisphere. Region-wise time series were cross-correlated using Spearman’s rank correlation to yield a subject-specific functional connectivity matrix (360×360), followed by Fisher-r-to-z transformations.

Diffusion MRI data were denoised and corrected for susceptibility distortions, head motion, and eddy currents using MRtrix3 (http://www.mrtrix.org/) (*6*). Structural connectivity (SC) was constructed from preprocessed diffusion MRI data using MRtrix. First, anatomically constrained tractography was performed using tissue types (grey matter, white matter, cerebrospinal fluid) derived from each participant’s preprocessed T1-weighted MRI, which was registered to the native diffusion MRI space. Second, multi-tissue response functions were estimated, and constrained spherical deconvolution and intensity normalization were performed. Seeding from all white matter voxels, a tractogram with 40 million streamlines (maximum tract length = 250; fractional anisotropy cut-off = 0.06) was generated via a probabilistic approach. Third, spherical deconvolution-informed filtering of tractograms (SIFT2) was applied to reconstruct whole-brain streamlines weighted by cross-sectional multipliers (*14*). Lastly, the reconstructed streamlines were mapped to the HCP-MMP1.0 atlas to generate a subject-specific SC matrix (360×360), in which connection weights between brain regions were defined as weighted streamline count. Group-level SC matrix was computed using distance-dependent thresholding to preserve the distribution of within- and between-hemisphere connection lengths in individual subjects (*15*). Prior to averaging, subject-specific SC matrix was log-transformed to reduce connectivity strength variance.

Quantitative T1 (qT1) images were registered to native Freesurfer space using boundary-based registration (*16*). Then, 14 equivolumetric cortical surfaces between the inner white and outer pial surfaces were generated (*1*, *17*). qT1 intensity values were sampled along these intracortical surfaces, mapped to the fsLR-32k template surface, smoothed with a 10-mm FWHM Gaussian kernel, and parcellated into 360 brain regions. Lastly, region-wise intensity profiles were cross-correlated using partial correlations that controlled for the average intensity profile, yielding a subject-specific micro-structural profile covariance matrix (360×360).

---

**Supplementary Figure 1.**
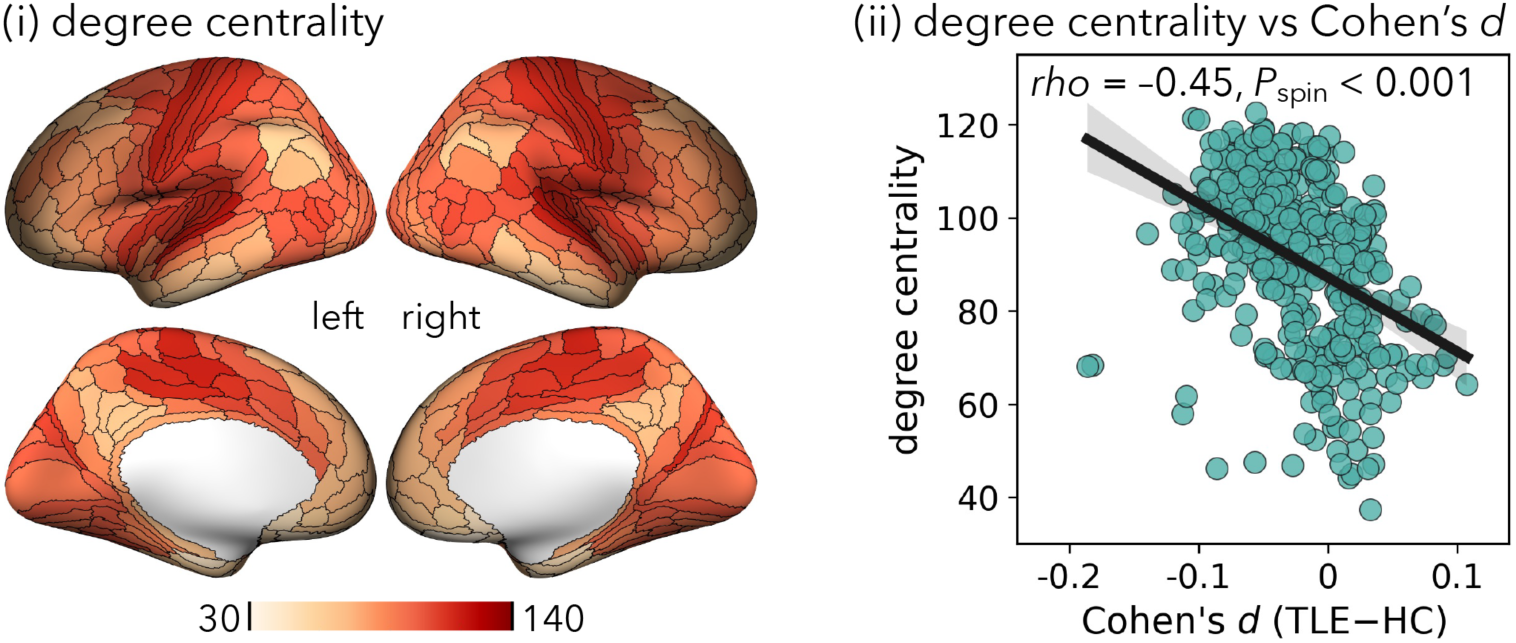
Relationship between regional connectivity strength and TLE FC alteration effect sizes. **(i)** Cortex-wide weighted degree centrality (node strength) derived from group-averaged FC matrix of healthy controls. **(ii)** Relationship between regional degree centrality and FC alteration effect size (Cohen’s *d* from Figure 2a) across the cortex. Each dot represents one region; the solid line denotes Spearman’s rank correlation with a 95% CI (shaded band). Significance (*P*_spin_) was assessed using spin permutation tests (5,000 rotations; one-sided).

**Supplementary Figure 2.**
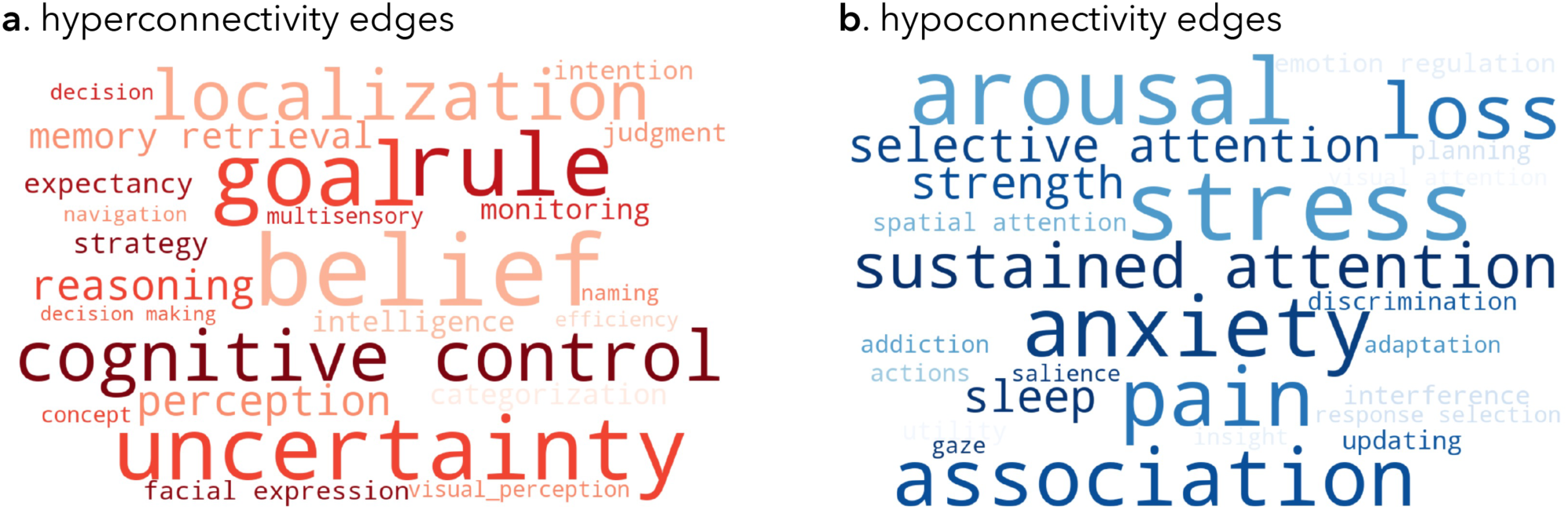
Neurosynth decoding of FC alteration patterns in TLE patients. Word clouds depict cognitive and behavioral terms associated with **(a)** hyperconnectivity and **(b)** hypoconnectivity patterns. Term relevance was quantified by the spatial similarity between each term’s Neurosynth meta-analytic activation map and FC effect size maps (from Figure 2b-c). Word size is proportional to the correlation coefficient (*rho*, 0.20–0.40), with larger words indicating stronger associations.

**Supplementary Figure 3.**
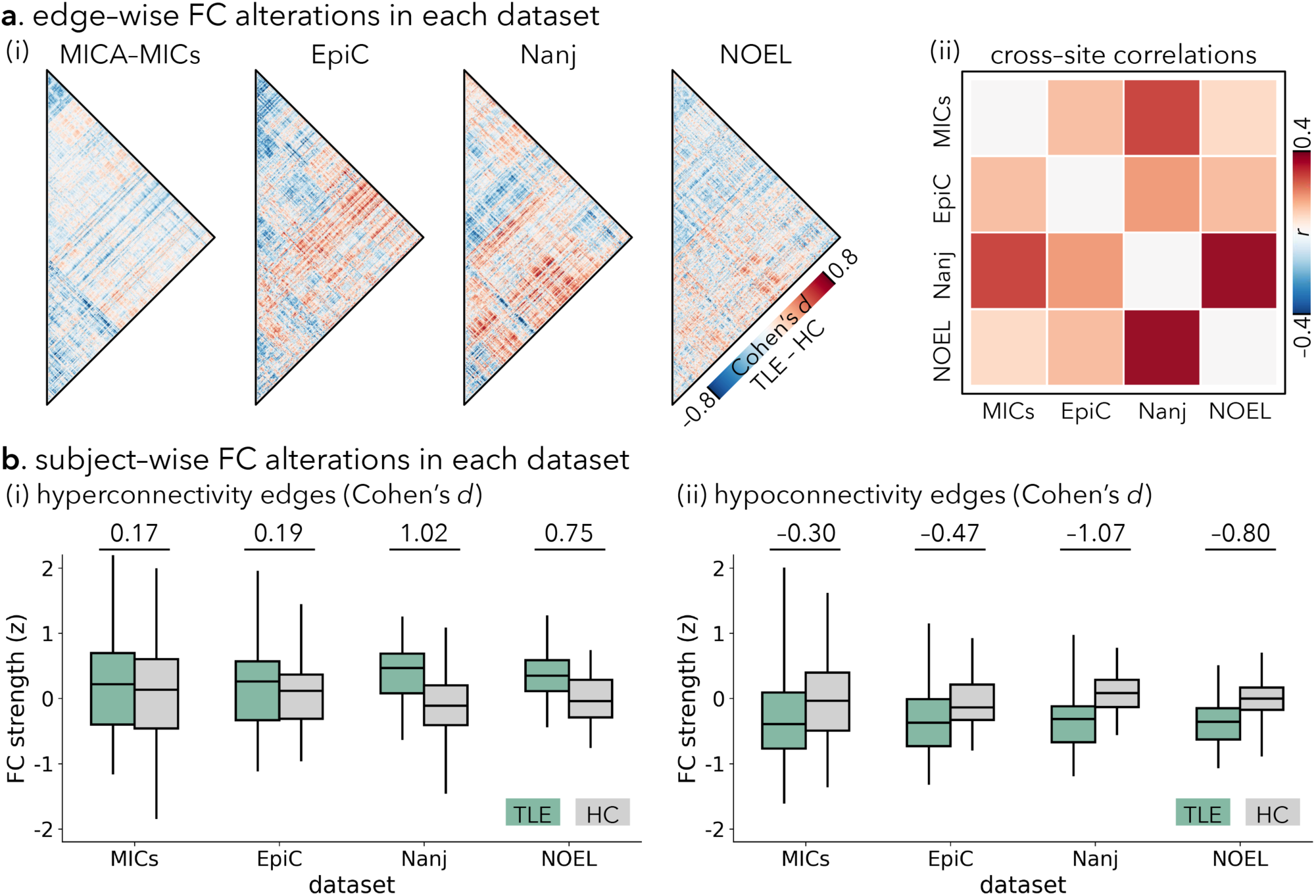
Cross-dataset consistency of FC alterations in TLE patients. **(a)** Dataset-specific effect size patterns. **(i)** Edge-wise TLE–control FC differences reported as Cohen’s *d* (warm, hyperconnectivity; cool, hypoconnectivity). **(ii)** Cross-dataset similarity of effect size patterns, shown as a correlation matrix of pairwise Pearson correlations between dataset-specific Cohen’s *d* vectors across all edges. **(b)** Within-dataset replication for edges identified in the full-sample analysis (see Figure 2b-c). Boxplots show individual mean FC values in TLE patients and healthy controls for edges exhibiting significant **(i)** hyperconnectivity and **(ii)** hypoconnectivity in the full-sample analysis. Boxplots show the median, interquartile range (25th−75th percentiles), and whiskers extending to 1.5× interquartile range.

**Supplementary Figure 4.**
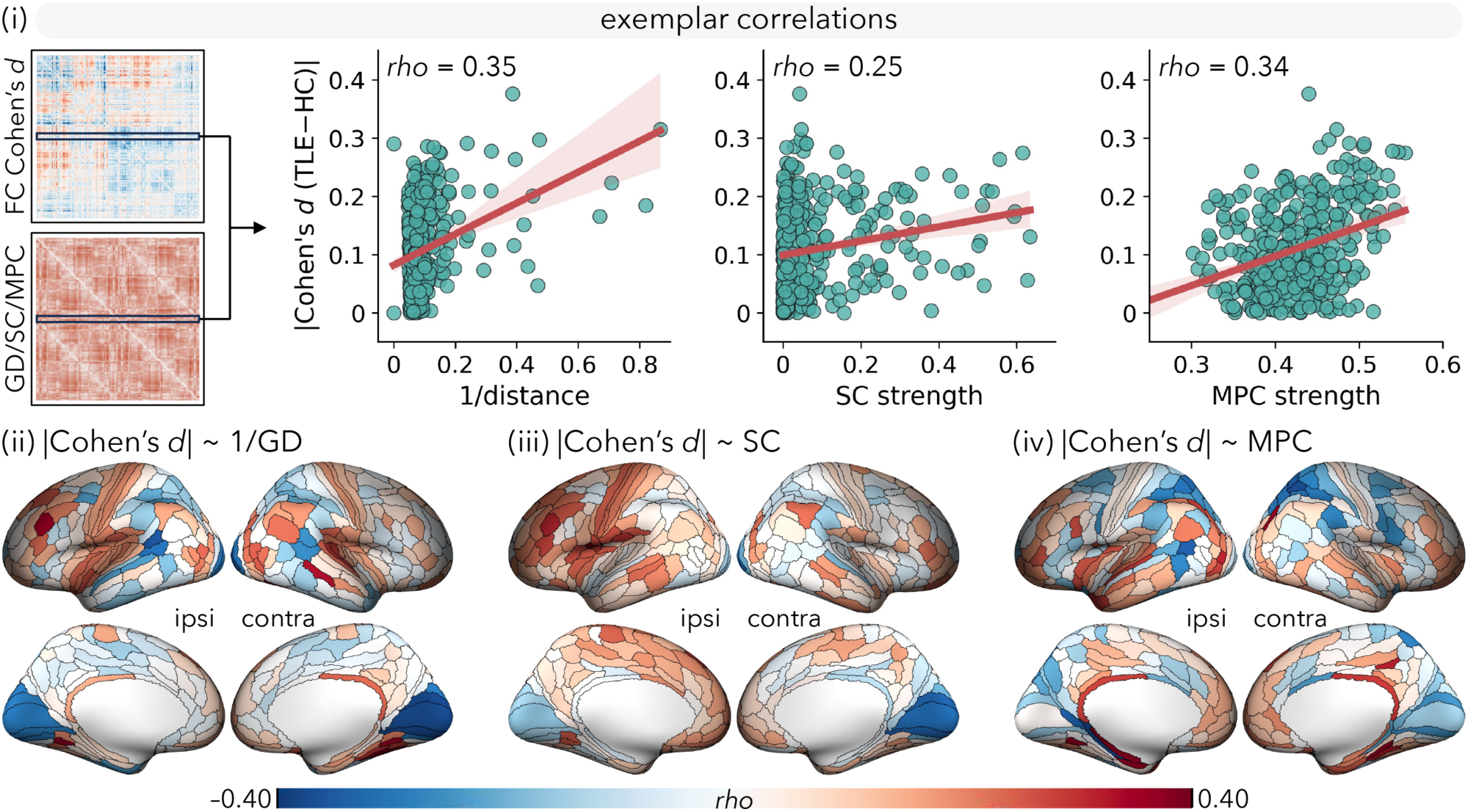
Region-wise associations between TLE-related FC alteration and structural affinity profiles. **(i)** Schematic example for a seed region showing the relationship between its FC alteration profile (Cohen’s *d* across edges to all other regions; from Figure 2a) and corresponding GD/SC/MPC profiles (see Figure 4a). Scatter plots show the Spearman’s rank correlations across regions; solid lines denote the fitted trend with a 95% CI (shaded band). **(ii–iv)** For each brain region, Spearman’s rank correlation (*rho*) was calculated between its Cohen’s *d* vector and the corresponding vector in **(ii)** GD, **(iii)** SC, and **(iv)** MPC. Region-wise *rho* values are projected onto the cortical surface (ipsi/contra indicate the hemisphere relative to the seizure focus). **Abbreviations**: ipsi, ipsilateral; contra, contralateral; GD, geodesic distance; SC, structural connectivity; MPC, microstructural profile covariance.

**Supplementary Figure 5.**
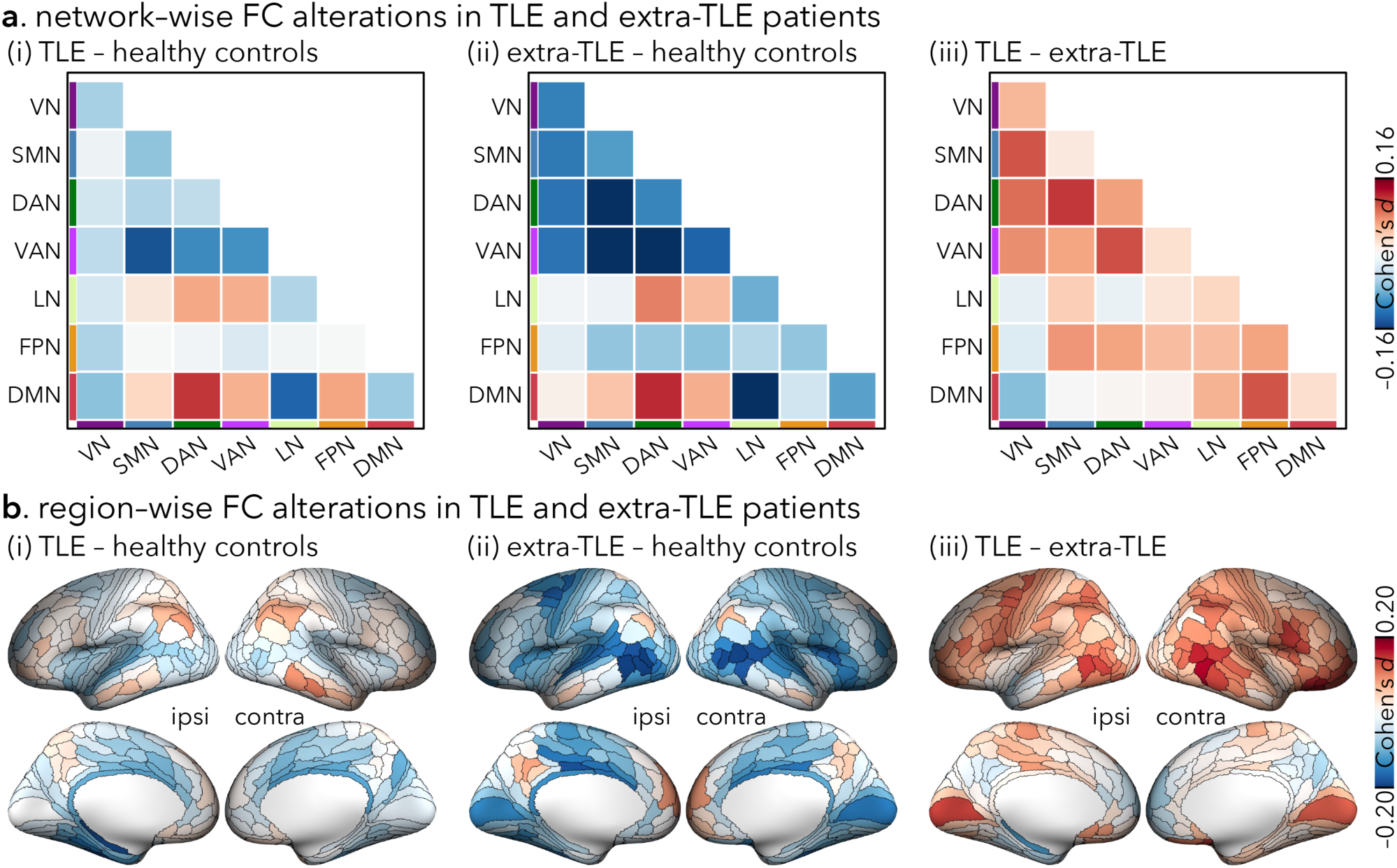
Disease specificity of FC alterations. **(a)** Network-level FC differences, quantified as network-wise Cohen’s *d* by averaging Cohen’s *d* across all edges within each functional network. **(b)** Region-level FC differences, quantified as region-wise Cohen’s *d* by averaging Cohen’s *d* across all edges incident on each brain region and projected onto the cortical surface (ipsi/contra indicate the hemisphere relative to the seizure focus). **(i-iii)** FC differences for **(i)** TLE patients versus healthy controls, **(ii)** extra-TLE patients versus healthy controls, and **(iii)** TLE patients versus extra-TLE patients. **Abbreviations**: ipsi, ipsilateral; contra, contralateral; VN, visual network; SMN, somatomotor network; DAN, dorsal attention network; VAN, ventral attention network; LN, limbic network; FPN, frontoparietal network; DMN, default mode network.

**Supplementary Figure 6.**
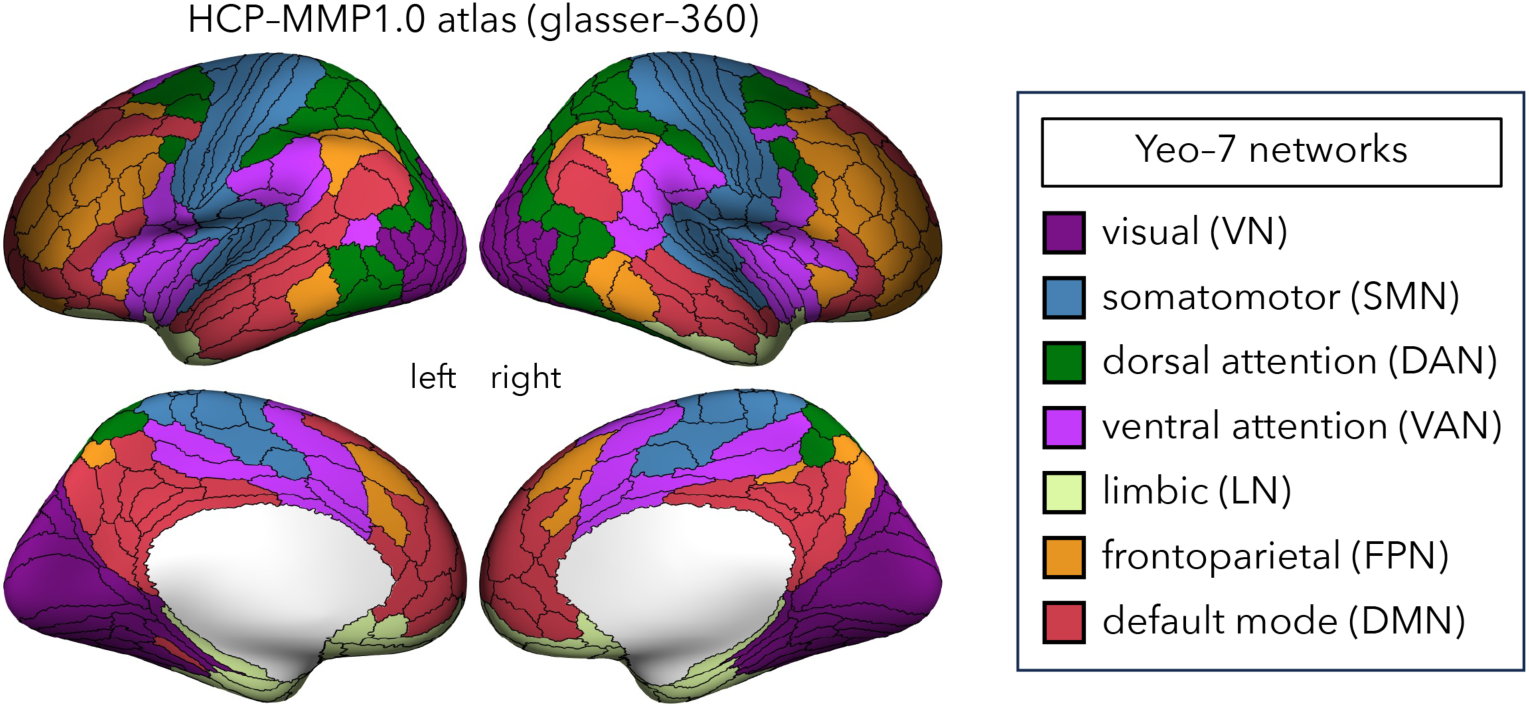
The HCP-MMP1.0 atlas with 360 regions at the surface. The cortical regions are assigned to each of the seven large-scale functional networks defined in the Yeo atlas, which include the visual (VN), somatomotor (SMN), dorsal attention (DAN), ventral attention (VAN), limbic (LN), frontoparietal (FPN), and default mode (DMN) networks.

